# Host gene-expression signatures accurately distinguish bacterial, viral, and inflammatory diseases in febrile children across multiple cohorts

**DOI:** 10.64898/2026.08.11.744042

**Authors:** Sandra Viz-Lasheras, Ana Dacosta, Irene Rivero-Calle, Federico Martinón-Torres, EUCLIDS, GENDRES, PERFORM, and DIAMONDS consortia, Alberto Gómez-Carballa, Antonio Salas

## Abstract

Accurate discrimination between viral, bacterial, and inflammatory diseases in febrile children remains a major clinical challenge that contributes to diagnostic uncertainty, inappropriate antimicrobial use, and suboptimal clinical management. Host blood transcriptomics offer a promising strategy to improve diagnostic precision. The present study represents the largest integrative multi-cohort pediatric study of transcriptomic biomarker discovery, validation, and confirmation reported to date, integrating harmonized public transcriptomic datasets with an independent confirmation cohort comprising well-phenotyped patients to identify parsimonious host-response signatures for differentiating viral, bacterial, and inflammatory diseases. Transcriptomic signatures were derived from an integrated retrospective microarray multi-cohort (*n*=1,683), independently validated in a retrospective RNA-seq cohort (*n*=767), and confirmed by digital PCR in an independent cohort (*n*=29), demonstrating reproducibility across patient populations, transcriptomic technologies, and analytical platforms. The analysis identified binary signatures and a unified multiclass classifier that consistently achieved high diagnostic accuracy across all three study phases and outperformed more than 30 published host transcriptomic signatures. Decision curve analysis showed substantially greater clinical net benefit than C-reactive protein across clinically relevant decision thresholds. These findings provide a strong foundation for clinically deployable molecular diagnostics to improve patient triage, antimicrobial stewardship, and precision medicine in childhood infections.

## Introduction

Infectious diseases continue to represent a major global health burden, being responsible for most deaths in children younger than five. Diagnosis remains challenging because clinical manifestations of bacterial and viral infections often overlap with those of non-infectious inflammatory conditions, hindering early diagnosis and appropriate treatment ^1^. This challenge is particularly pronounced in pediatrics, where nonspecific symptoms complicate identification of the underlying etiology. Microbiological confirmation by blood culture is typically delayed by 24–48 hours, limiting its value for early decision-making. Although molecular diagnostic techniques have improved pathogen detection, their clinical utility remains limited ^2^. Bacterial pathogens are often undetected because of low circulating loads or sampling difficulties. In contrast, viral detection in respiratory samples does not reliably establish causality, as similar carriage rates occur in symptomatic and healthy children ^3^. Improving etiological diagnosis is essential to optimize treatment, reduce unnecessary antibiotic use, and combat antimicrobial resistance ^4, 5^.

An alternative approach is the characterization of the host transcriptomic response to infection, which captures immune-mediated molecular changes following pathogen exposure. Advances in high-throughput transcriptomics and computational analysis now enable large-scale integration of heterogeneous datasets, facilitating pattern recognition and improving diagnostic and prognostic assessment ^6^. Host-response profiling is particularly valuable when direct pathogen detection is inconclusive or unavailable and has shown promise for distinguishing infectious etiologies, predicting disease severity, and identifying patients at risk of adverse outcomes. These advances have led to numerous host-response RNA signatures capable of distinguishing viral from bacterial infections and, in some cases, specific pathogens ^7–34^. However, existing signatures differ substantially in discovery cohorts, complexity, gene composition, validation strategies, and diagnostic performance. While some rely on large gene panels ^18, 23, 25, 26, 28, 34^, others use minimal signatures more suitable for rapid diagnostic devices ^7–11^. However, their reproducibility across populations, age groups, and transcriptomic platforms remains insufficiently evaluated.

The integration of transcriptomic data from multiple independent cohorts has emerged as a powerful strategy to overcome these limitations by increasing statistical power, reducing cohort-specific biases and overfitting, and enabling the identification of robust host-response biomarkers ^35^ that are reproducible across diverse populations, clinical settings, and platforms ^10, 36, 37^.

Accordingly, the present study reports the largest harmonized analysis of publicly available pediatric blood transcriptomic datasets to derive, validate, and benchmark compact blood-based gene expression signatures for the differential diagnosis of viral, bacterial, and non-infectious inflammatory diseases. The signatures were evaluated across independent cohorts and transcriptomic technologies, and systematically compared with more than 30 previously published host-based RNA signatures. The derived signatures demonstrate superior diagnostic accuracy, cross-platform reproducibility, and clinical transferability, providing a robust framework for the development of clinically deployable molecular diagnostics for pediatric febrile illness.

## Methods

### The discovery cohort

To develop novel transcriptomic signatures capable of distinguishing viral (VIR), bacterial (BAC), and inflammatory (INF) conditions in pediatric patients, a systematic search was conducted according to the PRISMA (Preferred Reporting Items for Systematic Reviews and Meta-Analyses) guidelines (**Figure S1A**). The Gene Expression Omnibus (GEO) repository was queried for human gene expression microarray datasets using the following terms: (“infection” OR “bacterial” OR “viral” OR “inflammatory”) AND (“pediatrics” OR “paediatrics” OR “children” OR “infants”) AND “blood” AND “Homo sapiens” [Organism] AND (GPL6947 OR GPL10558) AND “Expression profiling by array”). This search identified 36 eligible studies containing whole-blood Illumina BeadChip expression profiles from pediatric patients (<18 years) with acute febrile illnesses of viral, bacterial, or inflammatory origin, together with healthy controls (HC) (**Figure S1A**). After applying predefined inclusion and exclusion criteria, 14 datasets were retained (**Figure S1A**, **Table S1**, **Figure 1**). Samples with uncertain, mixed, or unclassified infectious or inflammatory etiologies were excluded from downstream analyses. Detailed cohort descriptions are provided in the original publications and datasets.

**Figure 1.**
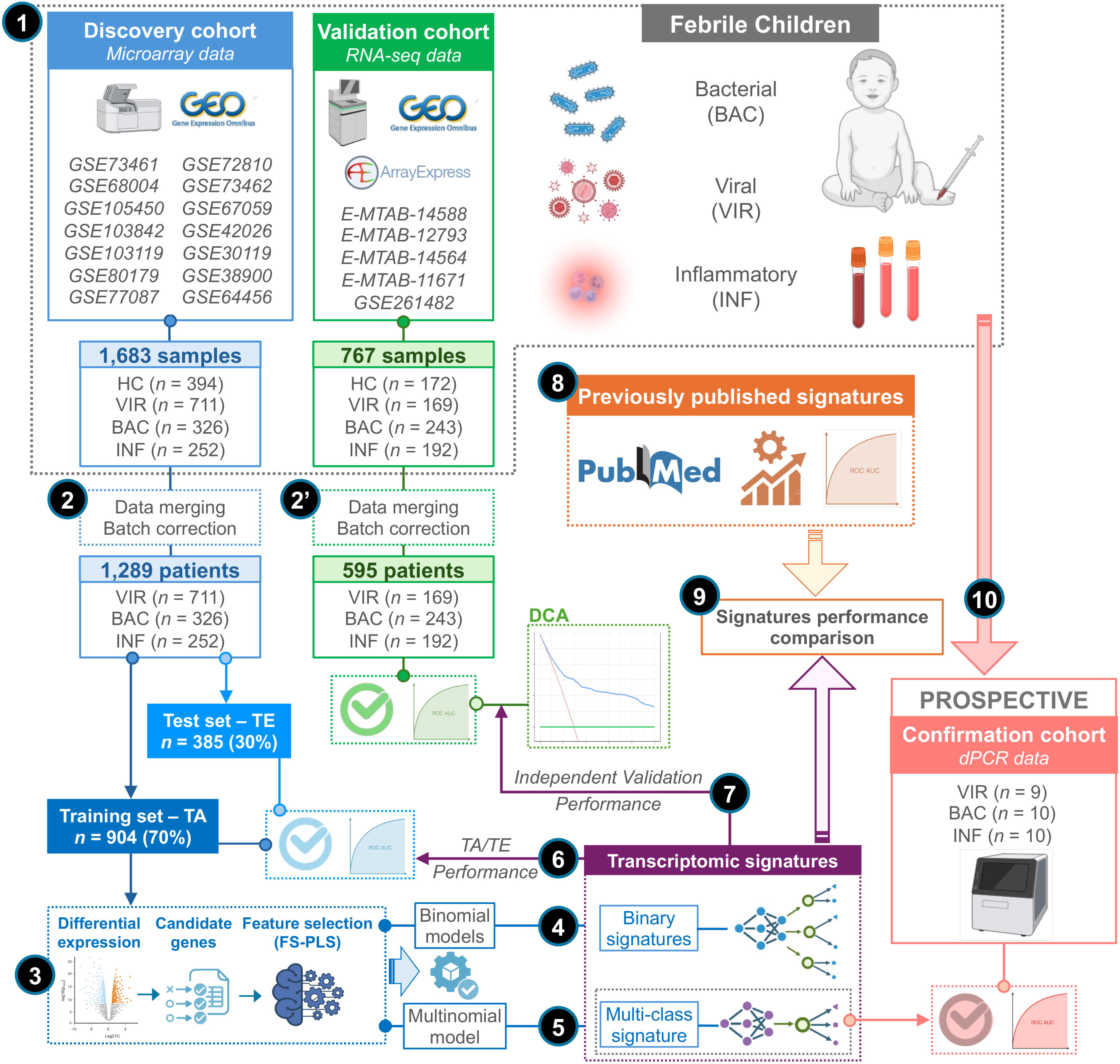
Overview of the study design. VIR, viral infections; BAC, bacterial infections; INF, inflammatory diseases. Step 1: Systematic identification, selection, and quality filtering of publicly available pediatric transcriptomic datasets to generate the discovery and validation cohorts. Step 2: Merging and batch correction of microarray datasets using *COCONUT*. Step 2′: Merging and batch correction of RNA-seq datasets using *ComBat-Seq*. Step 3: Differential gene expression analysis and feature selection to identify candidate biomarkers and derive transcriptomic signatures using FS-PLS. Step 4: Development of binary classification models. Step 5: Development of the multiclass classification model. Step 6: Evaluation of signature performance in the discovery cohort using the training and test sets. Step 7: Independent validation of the derived signatures in the RNA-seq cohort. Step 8: Systematic identification and selection of previously published host transcriptomic signatures. Step 9: Head-to-head benchmarking of newly derived and previously published signatures in the discovery and validation cohorts. Step 10: Recruitment of an independent pediatric confirmation cohort and validation of the multiclass signature by digital PCR (dPCR).

To integrate the selected studies, individual datasets were preprocessed and normalized using the *limma* package ^38^ and annotated using the *illuminaHumanv4.db* ^39^ or *illuminaHumanv3*.*db* ^40^ packages (version 1.26.0), depending on the microarray platform. When necessary, intra-study batch effects were first corrected using the *removeBatchEffect* function from *limma*. Subsequently, datasets were harmonized with *COCONUT* (COmbat CO-Normalization Using conTrols), which uses control samples to generate a single combined expression matrix while minimizing inter-study batch effects ^14, 37^ .After batch correction, HC samples were removed from subsequent analyses (**Figure 1**).

### The validation cohort

The diagnostic accuracy of the new RNA signatures was tested in an independent validation meta-cohort of public RNA-seq datasets from whole blood samples of pediatric febrile patients with VIR, BAC, and INF conditions, as well as HC. These publicly available datasets were generated within the framework of a European-funded consortium: DIAMONDS (Diagnosis and Management of Febrile Illness using RNA Personalised Molecular Signature Diagnosis; https://www.diamonds2020.eu), PERFORM (Personalised Risk Assessment in Febrile Illness to Optimise Real-life Management across the European Union; https://www.perform2020.org/), and EUCLIDS (European Union Childhood Life-Threatening Infectious Disease Study; https://www.diamonds2020.eu/our-research-history/euclids/). The corresponding transcriptomic datasets are available in ArrayExpress and GEO under the accession numbers E-MTAB-14588, E-MTAB-12793, E-MTAB-14564, E-MTAB-11671, and GSE261482 (**Table S1**, **Figure 1**).

Raw count matrices from the different sequencing batches were merged using the set of genes shared across all datasets. Batch effects between studies were corrected using the *Combat_seq* function from the *sva* R package ^41^. For visualization and other downstream analyses, normalized expression values were adjusted for age and sex using the *DESeq2* package ^38, 42^.

### Signatures discovery

To facilitate future clinical implementation, signature size was treated as a primary design criterion alongside diagnostic performance. Compact signatures (≤15–20 transcripts) were therefore prioritized to maximize their suitability for rapid point-of-care (POC) molecular diagnostics.

Signature discovery was performed using the harmonized discovery cohort, which was randomly divided into a 70% training (TA) and 30% test (TE) set using the *caret* R package ^43^, while preserving the original phenotype proportions (**Figure 1**). For determining the candidate genes that will be used as Forward Selection Partial Least Squares (FS-PLS) model input, differential expression analysis was carried out on the TA for the following phenotype comparisons: *i*) BAC *vs*. REST (VIR+INF), *ii*) VIR *vs*. REST (BAC+INF), and *iii*) INF *vs*. REST (BAC+VIR), using the *limma* package ^38^. A False Discovery Rate (FDR) threshold of 5% was applied. DEGs with an *P*_adj_<0.05, an |log_2_FC|>1 (FC: Fold Change), and a minimum average log_2_ expression of 4 were selected.

Feature selection was performed with Forward Selection Partial Least Squares (FS-PLS) ^44, 45^. Binary signatures were derived using multiple pairwise comparisons: VIR *vs*. REST (BAC+INF), VIR *vs*. BAC, VIR *vs*. INF; BAC *vs*. REST (VIR+INF), BAC *vs*. VIR, BAC *vs*. INF; and INF *vs*. REST (BAC+VIR), INF *vs*. VIR, INF *vs*. BAC. FS-PLS was also implemented within a multinomial framework (BAC *vs*. VIR, BAC *vs*. INF, VIR *vs*. INF) to derive a single multiclass signature capable of simultaneously distinguishing the three categories.

Model fitting was restricted to a maximum of 10-transcripts, and internal performance was evaluated using 10 repeated random subsampling cross-validations to assess classification stability across different data partitions. The selected biomarkers were then incorporated into generalized linear models (GLMs) using the *caret* package ^43^. Model coefficients estimated in the TA set were subsequently applied to generate predictions in the TE dataset.

### Predictive performance assessment

The predictive performance of both previously published and newly derived signature models was assessed by calculating the area under the receiver operating characteristic curve (AUC) and corresponding 95% confidence intervals (CI) using the *pROC* package ^46^. Optimal classification cut-points maximizing sensitivity/specificity were determined using the *OptimalCutPoints* package ^47^.

### The confirmation cohort

To confirm the performance of the multiclass signature in a clinically relevant setting and on an independent analytical platform, an independent confirmation cohort comprising 29 well-phenotyped pediatric patients selected from a prospectively established biobank was analyzed using digital PCR (dPCR). Clinical phenotypes were assigned according to the diagnostic algorithm developed by Herberg et al. ^7^.

The cohort included all clinical categories: 9 VIR, 10 BAC, and 10 INF cases, the latter comprising juvenile idiopathic arthritis (JIA; *n*=3), Kawasaki disease (KD; *n*=5), and Henoch–Schönlein purpura (HSP; *n*=2). The confirmation cohort had a mean age of 71.9 months and included 17 females (58.6%) and 12 males (41.4%).

Gene expression was quantified by dPCR using 10 pre-designed TaqMan assays targeting the selected transcripts (**Table S2**) and the Absolute Q™ Universal DNA Digital PCR Master Mix.

To minimize overfitting and improve model generalizability, a penalized multinomial logistic regression model was implemented using the *glmnet* R package ^48^. Model hyperparameters were optimized by repeated five-fold cross-validation (20 repeats) with automatic parameter tuning (*tuneLength*=10). The final model was selected according to the highest cross-validated accuracy, and the resulting class probabilities were used to evaluate diagnostic performance.

To assess whether the diagnostic signature could be further simplified while retaining its discriminatory performance, an additional multinomial logistic regression model was retrained after excluding selected markers. Model training, hyperparameter optimization, and performance evaluation were performed following the same procedure described above.

### Clinical utility of the predictive model

To further evaluate the clinical utility of the proposed signatures, Decision Curve Analysis (DCA) was performed in the validation cohort using the *rmda* R package ^49^. Unlike conventional performance metrics (e.g., AUC, sensitivity, and specificity), DCA estimates the net clinical benefit of a diagnostic model across a range of decision thresholds, providing an assessment of its potential value for clinical decision-making. The multiclass transcriptomic signature was compared with standard management strategies (“treat all” and “treat none”) and CRP-based decision thresholds (≥60 mg/L for bacterial infection and <60 mg/L for viral infection).

### Cross-comparison of transcriptomic signatures

To identify previously published host transcriptomic signatures for differentiating viral and bacterial infections, a systematic PubMed search was conducted according to PRISMA guidelines using the following query: (“bacterial”[Title/Abstract] OR “viral”[Title/Abstract]) AND (“host”[Title/Abstract] OR transcript*[Title/Abstract] OR “gene expression”[Title/Abstract] OR RNA[Title/Abstract]) AND (signature[Title/Abstract] OR classifier[Title/Abstract]) AND (blood[Title/Abstract] OR peripheral[Title/Abstract]) AND “Humans”[Mesh]. The search retrieved 303 records, with 9 additional studies identified through manual screening of reference lists and relevant reviews, yielding 312 records (**Figure S1B**). Only studies with publicly available transcriptomic data were included. The recently published *Mycoplasma pneumoniae* signature ^27^ was excluded because it was specifically designed to detect *M. pneumoniae* infection and therefore is not applicable to the broader diagnostic categories evaluated here. After applying the predefined selection criteria (**Figure S1B**), 32 transcriptomic signatures from 28 publications were retained (**Table S3**) ^7–34^. Only transcripts whose mapped gene symbols were present in both the discovery and validation datasets were retained for downstream analyses (**Table S3**). Detailed cohort information is available in the original publications.

To enable a fair comparison with the newly derived signatures, all published models were retrained using the unified analytical framework developed in the present study. Model coefficients were re-estimated from the TA microarray dataset using binomial generalized linear models (*glm*; *caret* R package ^50^) and subsequently evaluated, without further parameter tuning, in the TE dataset, the combined discovery cohort (TA+TE), and the independent RNA-seq validation cohort. Performance metrics were calculated using the same evaluation pipeline applied to the newly developed signatures.

## Results

### Discovery and validation cohorts

Following systematic curation of GEO datasets according to the predefined inclusion criteria (**Figure S1A**), 14 pediatric microarray cohorts comprising patients with viral, bacterial, or inflammatory diseases, together with healthy controls (HCs), were integrated into the discovery dataset. After dataset-specific normalization, principal component analysis (PCA) of individual studies showed a clear separation by phenotypic group (BAC, VIR, INF, and HC), indicating that the transcriptomic profiles captured biologically meaningful differences among clinical conditions (**Figure S2**).

The harmonized discovery dataset comprised 1,683 samples, including 711 VIR, 326 BAC, 252 INF (66 juvenile idiopathic arthritis [JIA], 168 Kawasaki disease [KD], and 18 Henoch–Schönlein purpura [HSP]), and 394 HCs (**Figure 1**, **Table S1**). Before batch correction, PCA revealed clustering by study rather than phenotype (**Figure S3**). Subsequently, COCONUT harmonization effectively removed inter-study variation while preserving biological separation between disease groups (**Figure S3**). The integrated cohort was randomly divided into a training (TA; 70%, *n*=904) and test (TE; 30%, *n*=385) set. The TA cohort included 498 VIR, 229 BAC, and 177 INF cases (45 JIA, 119 KD, and 13 HSP), whereas the TE cohort comprised 213 VIR, 97 BAC, and 75 INF cases (21 JIA, 49 KD, and 5 HSP).

The validation cohort consisted of five independent RNA-seq datasets (**Figure 1**, **Table S3**), comprising 767 pediatric transcriptomes: 169 VIR, 234 BAC, 192 INF (50 JIA and 142 KD), and 172 HCs. The mean age was 63.9 months, with 346 females (42.5%) and 468 males (57.5%). For age-stratified analyses, participants were grouped into 0–3, >3–6, >6–24, and >24 months. Following ComBat-Seq correction and normalization, PCA confirmed successful harmonization, with no residual clustering by study (**Figure S4**). The two cohorts were demographically comparable, with statistically significant differences observed only in donor age within the viral category (α = 0.01; **Table S1**).

### Binary signatures in the discovery and validation cohorts

Differential expression analysis of the discovery cohort, followed by filtering (*P*_adj_<0.05, |log_2_FC|>1, and mean log_2_ expression >4), identified 53, 55, and 37 DEGs in the VIR *vs*. BAC+INF, BAC *vs*. VIR+INF, and INF *vs*. VIR+BAC comparisons, respectively (**Figure 2A**, **Table S4**). Upregulated transcripts predominated in all contrasts. Specifically, 35/18, 32/23, and 19/18 genes were up- and downregulated in the VIR, BAC, and INF comparisons, respectively (**Figure 2A**).

**Figure 2.**
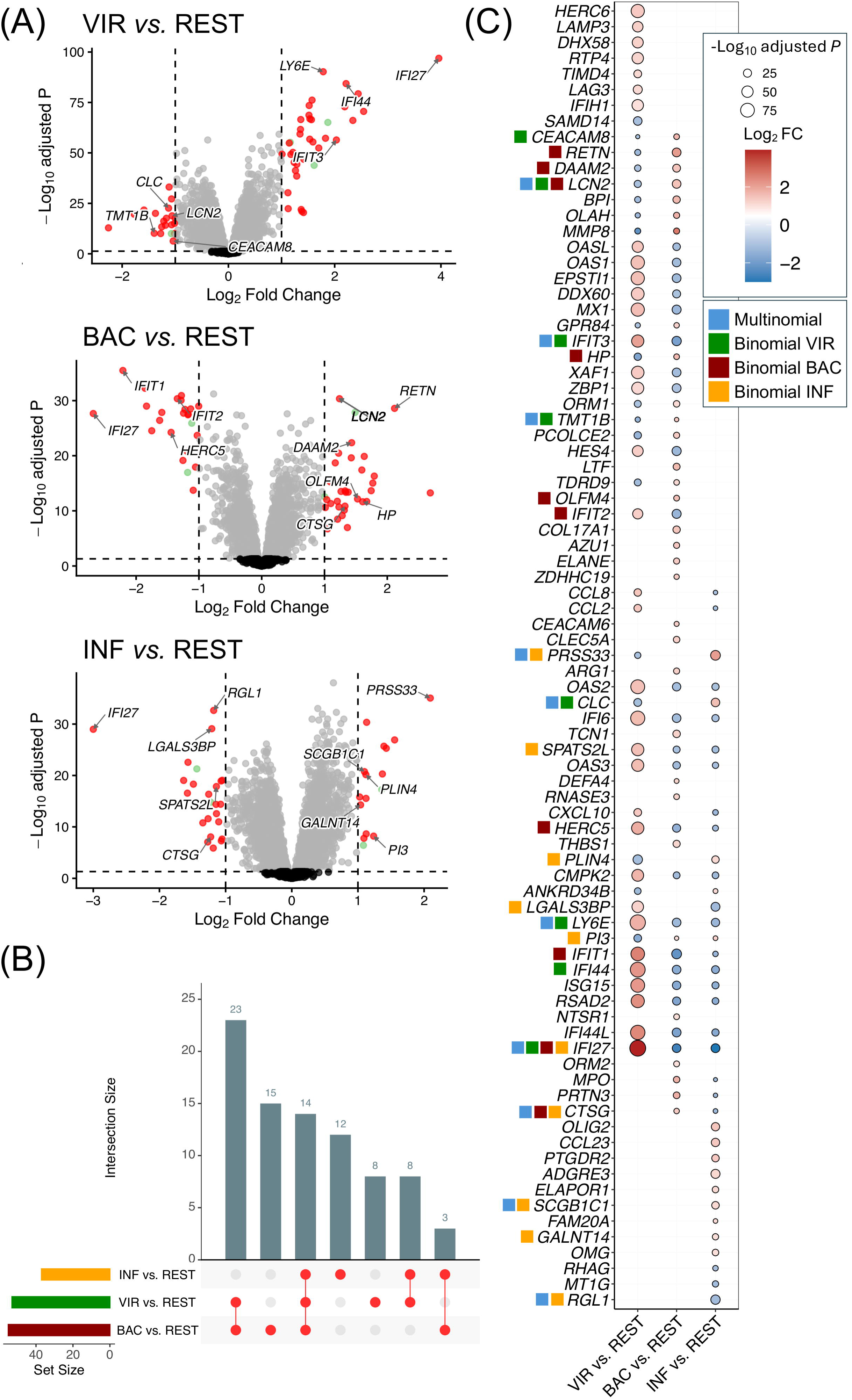
Differentially expressed genes (DEGs) between disease groups. A) Volcano plots showing the DEGs between conditions: VIR *vs.* REST, BAC *vs.* REST, and INF *vs.* REST. In red, the selected biomarker to develop the predictive models. Genes with an *P*_adj_< 0.05 and a |log_2_FC)>1 and an Averexp>4 were colored in red, genes with an *P*_adj_< 0.05 and a |log_2_FC)|>1 and an Averexp<4 were colored in light green, genes with an *P*_adj_< 0.05 but a |log_2_FC)|>1 were colored in grey and non-significant genes with |log_2_FC)|>1 were colored in black. B) Upset plot representing common and unique DEGs from each comparison. C) Dotplot of log_2_FC differences in the 3 comparisons (BAC *vs*. REST, VIR *vs.* REST, and INF *vs*. REST) for the candidate genes. Color scale indicates log_2_FC values, and the size of the dots represents the - log_10_ *P*_adj_. Genes included in the signatures are indicated by colored squares preceding the gene name.

Among all DEGs, 14 were shared across the three comparisons, whereas 23 were common to the VIR and BAC contrasts. In contrast, only three transcripts overlapped between BAC and INF conditions, highlighting the greater transcriptional similarity between infectious phenotypes and the distinct molecular profile of non-infectious inflammatory diseases (**Figure 2B-C**, **Table S4**).

A total of 83 unique DEGs were retained for feature selection using FS-PLS (**Figure 2C**). Three binary classifiers were subsequently derived to distinguish VIR, BAC, and INF diseases from all remaining conditions. The optimal models comprised 8 transcripts for the viral signature (*IFI27*, *LCN2*, *CLC*, *IFI44*, *TMT1B*, *IFIT3*, *CEACAM8*, *LY6E*), ten genes for the bacterial signature (*IFIT1*, *RETN*, *IFI27*, *HP*, *HERC5*, *LCN2*, *IFIT2*, *CTSG*, *OLFM4*, *DAAM2*) and ten genes for the inflammatory signature (*RGL1*, *PRSS33*, *PLIN4*, *CTSG*, *IFI27*, *GALNT14*, *SCGB1C1*, *LGALS3BP*, *SPATS2L*, *PI3*) (**Figure 2C**, **Table S5**).

The 8-transcript viral signature differentiated VIR *vs.* BAC+INF conditions with an AUC of 0.93 in both the TA and TE subsets (95% CI_TA_: 0.92–0.95; 95% CI_TE_: 0.90–0.95); **Figure 3A**. Bacterial signature discriminated BAC infection from other conditions (VIR+INF), yielding AUCs of 0.88 (95% CI_TA_: 0.86–0.91) in the TA and 0.91 (95% CI_TE_: 0.88–0.95) in the TE (**Figure 3B**). The inflammatory signature accurately distinguished INF diseases from VIR and BAC conditions, showing an AUC of 0.95 (95% CI_TA_: 0.93–0.97) in the TA and 0.94 (95% CI_TE_: 0.92–0.96) in the TE (**Figure 3C**).

**Figure 3.**
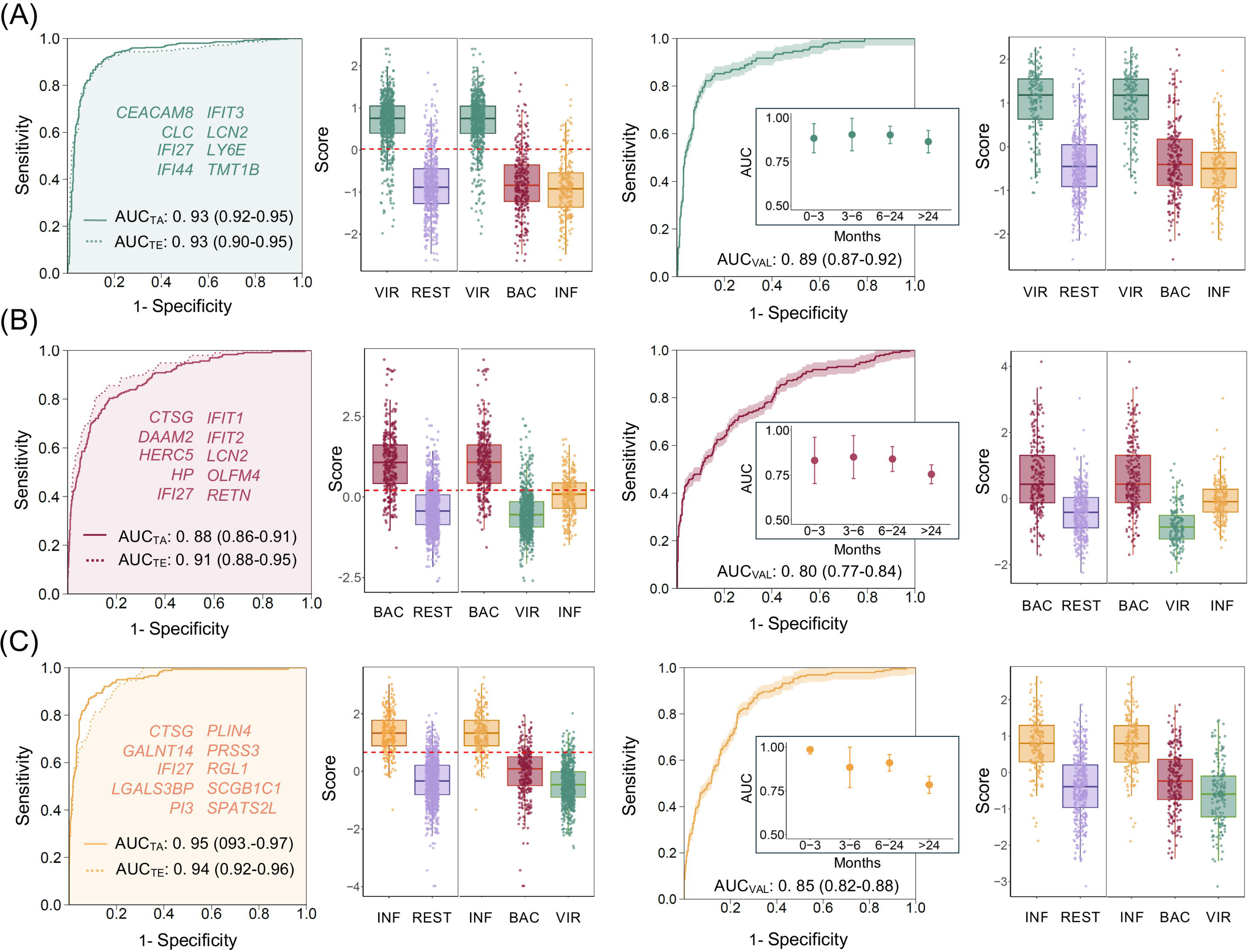
Receiver operating characteristic (ROC) curves and area under the curve (AUC) values for the three binary classifiers: viral (A), bacterial (B), and inflammatory (C), evaluated in the discovery cohort (training [TR] and test [TE] sets) and the independent validation cohort (VAL). Boxplots show the distribution of predicted probabilities for each disease group, with the red dashed line indicating the optimal classification threshold. Insets display AUC values stratified by age group in the validation cohort.

External validation in the independent RNA-seq meta-cohort confirmed the robustness of the three signatures, with AUCs of 0.89 (95% CI: 0.87–0.92) for VIR *vs*. REST, 0.80 (95% CI: 0.77–0.84) for BAC *vs*. REST, and 0.85 (95% CI: 0.82–0.88) for INF *vs*. REST (**Figure 3**). Consistently, boxplots of the per-sample scores derived from the diagnostic binary models demonstrated statistically significant discrimination between each target category (VIR, BAC, and INF) and the corresponding non-target samples across all comparisons in both the discovery and validation datasets. Performance remained consistent across all pediatric age groups (0–3, >3–6, >6–24, and >24 months). The viral signature achieved AUCs of 0.88, 0.90, 0.90, and 0.86, respectively. The bacterial signature remained stable (0.83–0.85 in children ≤6 months, 0.84 in >6–24 months, and 0.76 in >24 months), while the inflammatory signature achieved AUCs of 0.98, 0.85, 0.91, and 0.80, respectively (**Figure 3**).

### Discovery and validation of the multi-class signature

A multinomial model was developed to derive a unique transcriptomic signature capable of differentiating among the three disease phenotypes, thus simplifying the use and interpretation of the predictive test in more complex disease scenarios. The same 83 candidate genes were used as input for the FS-PLS multinomial analysis. The multinomial model identified a ten transcripts-signature (*IFI27*, *LCN2*, *CLC*, *CTSG*, *RGL1*, *IFIT3*, *LY6E*, *SCGB1C1*, *TMT1B* and *PRSS33*) that accurately differentiate between VIR, BAC and INF infections with AUCs in the discovery cohort of 0.94 (TA) and 0.93 (TE) (95%CI_TA_: 0.92– 0.96; 95%CI_TE_: 0.91–0.96; ; **Figure 4A**) for VIR infections, 0.89 (TA) and 0.90 (TE) (95%CI_TA_: 0.86–0.92; 95%CI_TE_: 0.86–0.93; **Figure 4B**) for BAC infections and 0.95 (TA) and 0.94 (TE) (95%CI_TA_: 0.93–0.96; 95%CI_TE_: 0.92–0.97;**Figure 4C**) for the INF diseases. Among the genes selected by the multinomial model, *IFI27*, *CTSG*, *RGL1*, *SCGB1C1*, and *PRSS33* were also present in the binary classification signatures (**Figure 2C**).

**Figure 4.**
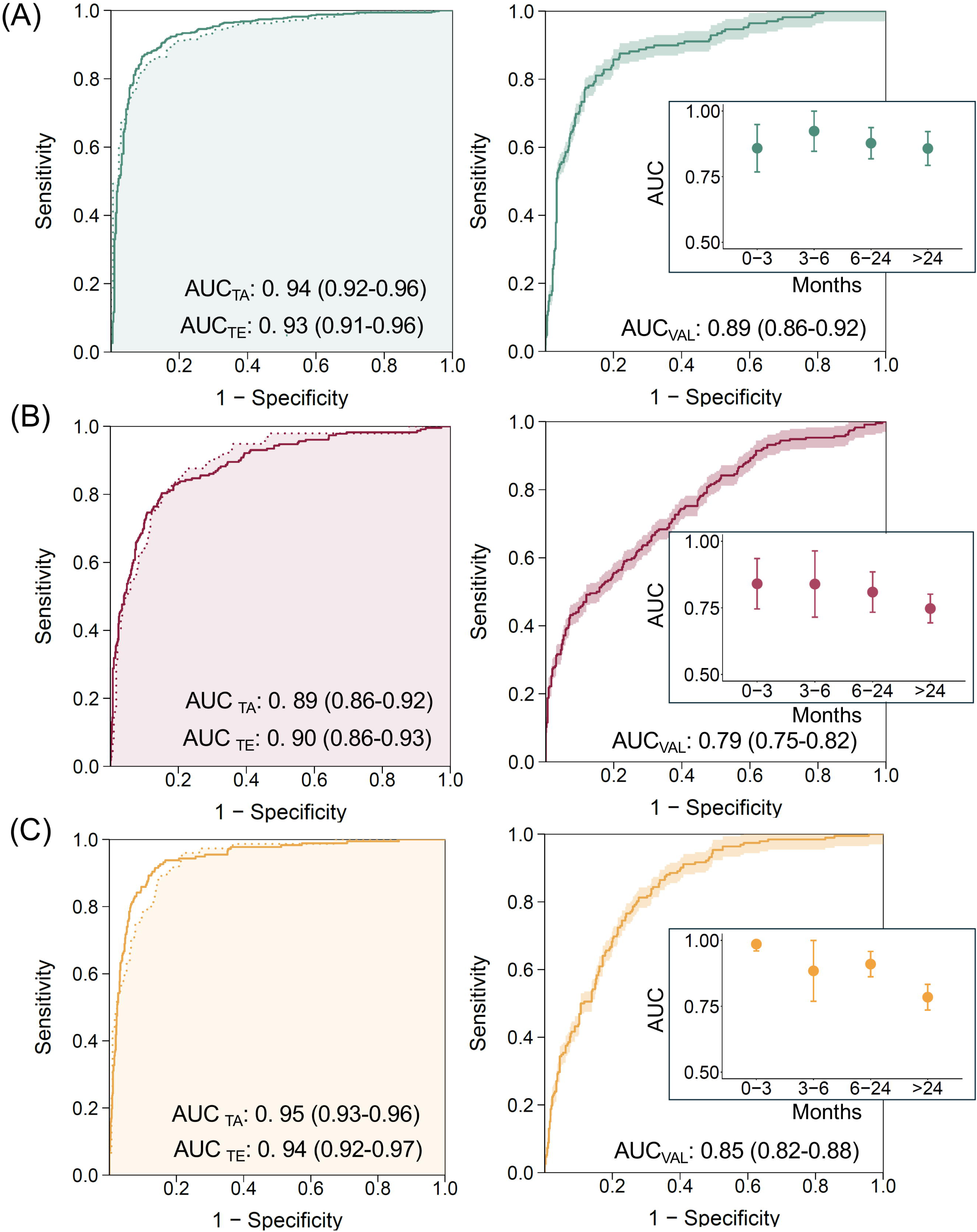
Receiver operating characteristic (ROC) curves and area under the curve (AUC) values for the multiclass classifier evaluated in the discovery cohort (training [TR] and test [TE] sets) and the independent validation cohort (VAL). ROC curves are shown for the one-versus-rest contrasts: (A) viral infections (VIR) *vs*. all other conditions, (B) bacterial infections (BAC) *vs*. all other conditions, and (C) inflammatory diseases (INF) *vs*. all other conditions. Insets show AUC values stratified by age group in the validation cohort.

The signature derived from the three multinomial model demonstrated a similar accuracy in differentiating the three phenotypes of interest in the RNA-seq validation cohort: AUC of 0.89 (95%CI: 0.86–0.92; **Figure 4A**) for VIR infections, 0.79 (95%CI: 0.75–0.82; **Figure 4B**) for BAC infections, and 0.85 (95%CI: 0.82–0.88) for INF conditions (**Figure 4C**). The performance of the multi-class signature in the validation cohort was fully comparable with that obtained with the binary signatures.

Consistent with the binary classifiers, the multiclass signature maintained high diagnostic accuracy across all clinically relevant age groups and disease categories in the validation cohort. For VIR infections, discrimination remained consistently strong, with AUC values of 0.86 in neonates (0–3 months), 0.92 in infants aged >3–6 months, 0.88 in children aged >6–24 months, and 0.86 in those older than 24 months (**Figure 4A**). Performance for BAC infections was slightly lower but remained stable across age groups, yielding AUCs of 0.84, 0.84, 0.81, and 0.75, respectively (**Figure 4B**). Likewise, the multiclass model accurately identified INF diseases across all age strata, achieving AUC values of 0.99 in neonates, 0.88 in infants aged >3–6 months, 0.91 in children aged >6–24 months, and 0.78 in those older than 24 months (**Figure 4C**).

### Decision curve analysis

Decision curve analysis (DCA) was performed to evaluate the clinical utility of the new signatures derived from the multi-cohort meta-analysis, adding contextual value over the traditional assessment previously described. DCA of the binomial classifiers demonstrates that the new transcriptomic signature (BAC, VIR, and INF) offers meaningful clinical utility across a broad range of decision thresholds. In the total cohort, all binomial models consistently achieved a higher net benefit than the default “treat-all” and “treat-none” strategies, indicating their potential value for phenotype-specific decision-making across clinically relevant threshold probabilities (**Figure 5A**). Specifically, the bacterial classifier exhibited sustained positive net benefit from low to very high threshold probabilities, with superiority over default strategies extending up to thresholds close to 0.95 (**Figure 5A**), supporting its robustness for guiding antibiotic treatment decisions under both permissive and conservative clinical scenarios. The viral classifier demonstrated a clear advantage over “treat-all” and “treat-none” strategies at low-to-moderate threshold probabilities (∼0.1–0.6), after which the net benefit declined, consistent with reduced clinical utility at very high thresholds where extreme certainty is required (**Figure 5A**). Similarly, the inflammatory classifier showed improved net benefit over default strategies across a wide threshold range, with gradual attenuation at higher thresholds (**Figure 5A**).

**Figure 5.**
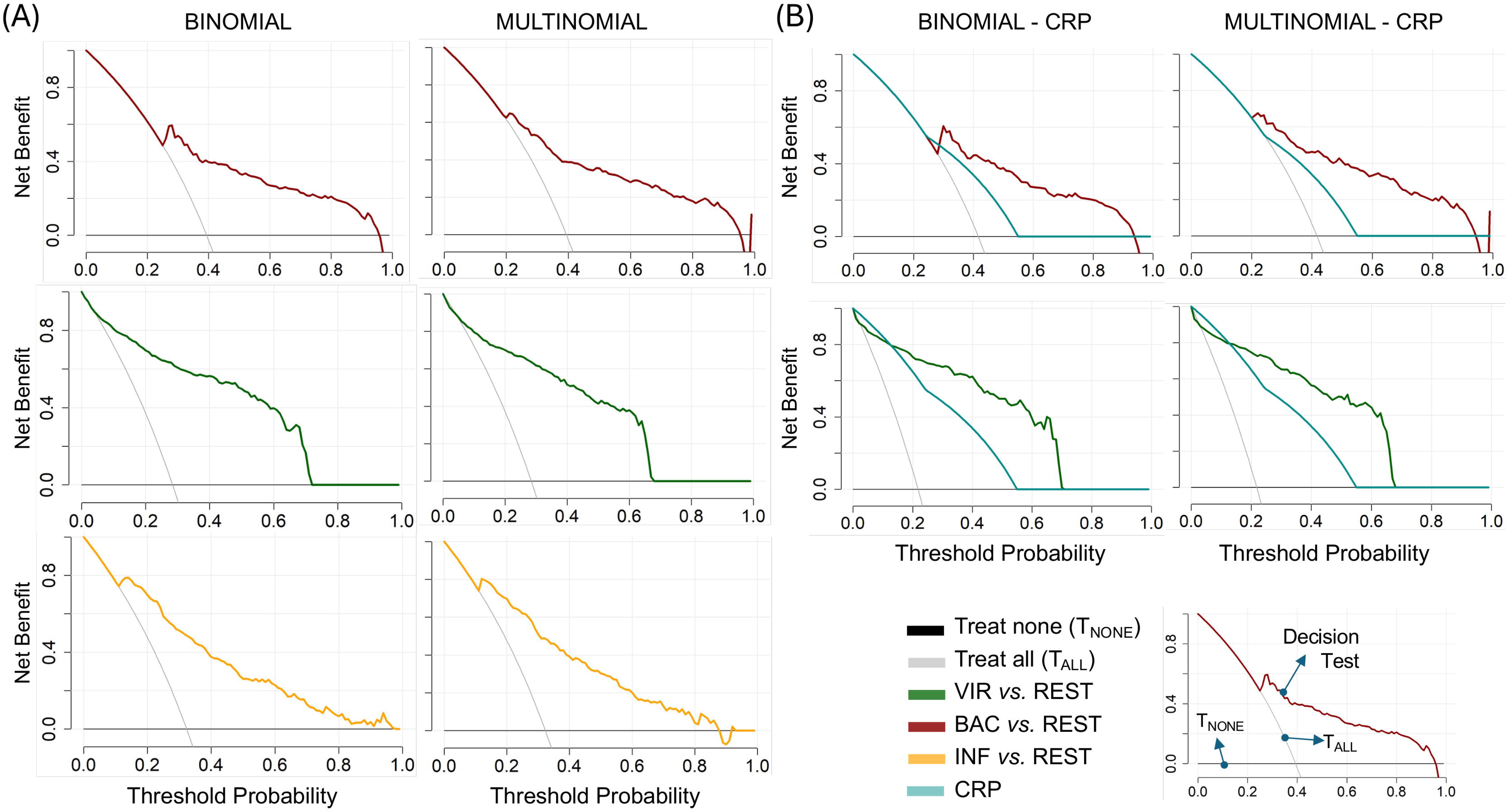
Decision curve analysis (DCA) evaluating the clinical utility of the transcriptomic classifiers. (A) Net benefit of the multiclass classifier in the entire validation cohort for the contrasts BAC *vs*. REST, VIR *vs*. REST, and INF *vs*. REST. (B) Net benefit of the binary classifiers in the subset of patients with available C-reactive protein (CRP) measurements, comparing transcriptomic signatures with CRP-based clinical decision thresholds for BAC *vs*. REST and VIR *vs*. REST.

Consistent with these findings, the multinomial transcriptomic model demonstrates a higher net benefit than both the “treat-all” and “treat-none” strategies across a broad spectrum of threshold probabilities, particularly for the classification of BAC infections. Notably, its performance was largely comparable to that observed for the corresponding binomial models (**Figure 5A**). Again, the multiclass signature yielded the highest net benefit for guiding antibiotic treatment in BAC infections at threshold probabilities above 0.2, extending up to threshold probabilities of 0.95 (**Figure 5A**). The patterns of DCA for the diagnosis of VIR and INF groups are also comparable to those observed in the binomial model (**Figure 5A**).

When restricting the analysis to patients with available CRP measurements (*n=*353; 146 BAC, 77 VIR, and 130 INF), the binomial transcriptomic signatures maintained their advantage over standard clinical approaches. In this subset, all three classifiers yielded a higher net benefit than CRP-guided strategies and the extreme treatment rules, reinforcing the added value of transcriptomic profiling beyond conventional inflammatory markers (**Figure 5B**). Similarly, for the transcriptomic multinomial model, greater net benefit than CRP and the standard strategies was observed at threshold probabilities between approximately 0.1 and 0.2, for diagnosis of BAC infections and INF conditions. This advantage persisted up to thresholds of around 0.7 in the case of the VIR group and above 0.95 for the BAC patients (**Figure 5B**). This improvement was particularly evident across clinically actionable threshold ranges (∼0.1–0.7 for viral decision-making and ∼0.3–0.95 for bacterial infection management), where CRP-based strategies showed diminishing or negligible net benefit (**Figure 5B**). Importantly, at threshold probabilities above 0.5 (corresponding to more conservative clinical decision-making), the net benefit of CRP rapidly diminishes, suggesting limited clinical utility in this range. In contrast, all transcriptomic signatures retain a meaningful positive net benefit, supporting the ability to inform decisions even when a high level of confidence in viral etiology is required (**Figure 5B**).

### Comparison of signatures performance

The performance of the new proposed multinomial (10-transcripts) and binomial (10- and 8-transcripts) signatures was systematically compared for the individual discrimination of VIR, BAC, and INF conditions against a broad panel (*n*=32) of previously published host-response transcriptional signatures spanning a wide range of sizes (2 to 398 transcripts) and methodological approaches (**Table S6**). As expected, larger signatures tend to achieve higher AUCs in the discovery cohort. Many of these large signatures exhibited substantial performance loss in the validation cohort, in some cases reaching a chance-level discrimination (e.g., 160-Habgood-Coote et al., 398-Andres-Terre et al., **Table S6**).

In the VIR *vs*. REST comparison, the 8-transcript signature achieved an AUC of 0.93 in the discovery cohort, placing it at the top of performing compact signatures, exceeding the performance of several widely cited small, published models (**Table S6; Figure 6**), including 4-Sampson et al. (AUC=0.84), 7-Sweeney et al. (AUC=0.90), and 4-Falsey et al. (AUC=0.87). Although a small number of very large signatures reached higher discovery AUCs, these models required between 29 and 398 transcripts (**Table S6; Figure 6**). In the validation cohort, the viral signature retained strong performance (AUC=0.89), being surpassed only by a limited number of compact published signatures, including 7-Sweeney et al. (AUC=0.91) and 8-Rao et al. (AUC=0.91), the latter having an equivalent gene set size.

**Figure 6.**
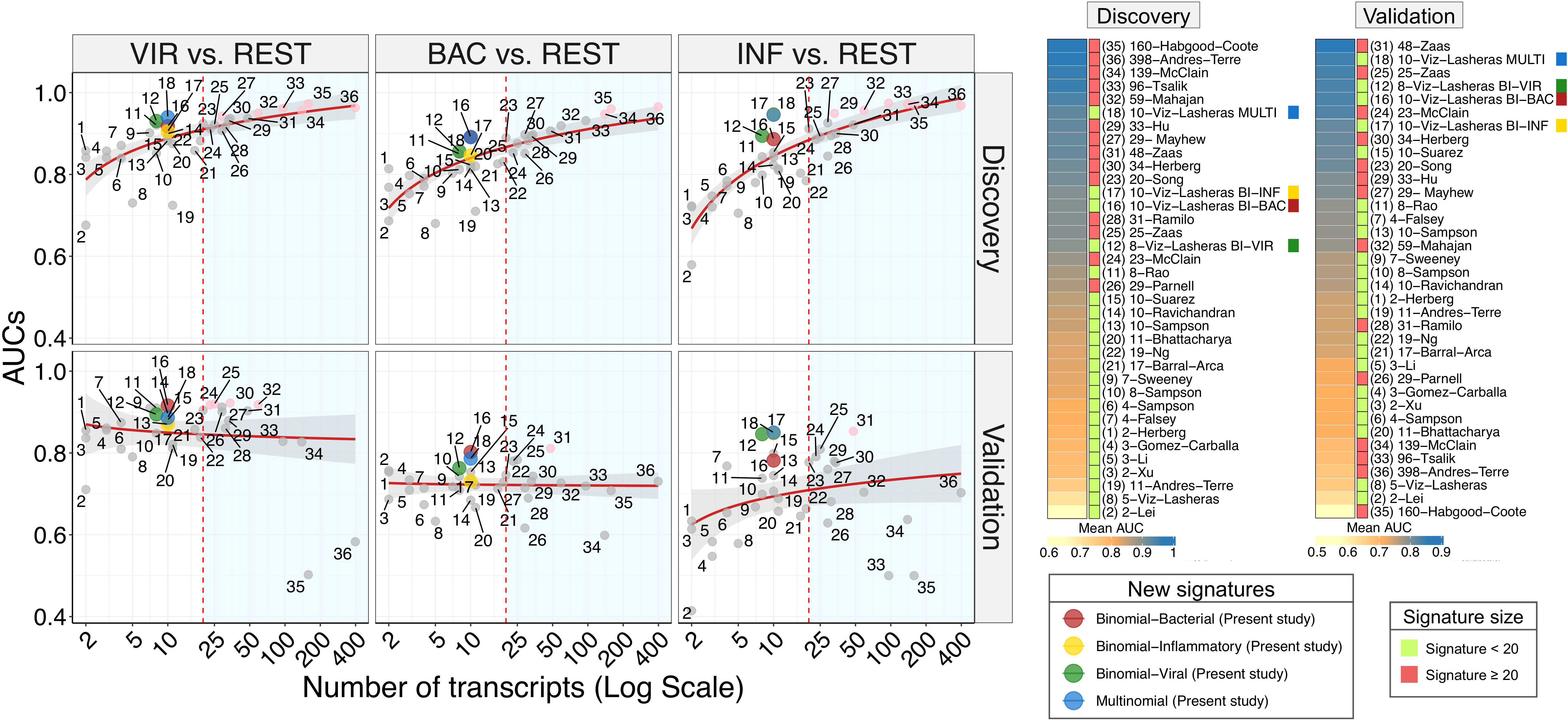
Relationship between signature size and diagnostic performance. Correlation between AUC values and the number of transcripts included in each evaluated signature, comprising both the newly derived binary and multiclass signatures and previously published transcriptomic signatures, assessed in the discovery and validation cohorts. The x-axis is displayed on a logarithmic scale. The red dashed vertical line indicates a signature size threshold of 20 transcripts.

A similar pattern was observed for bacterial classification, where the binary signature achieved AUCs of 0.88 in the discovery cohort, a value that exceeds all those of published signatures with similar or substantially smaller gene sets, and the maximum AUC value in the validation cohort (AUC=0.80); **Figure 6**. Similarly, the inflammatory binary signature showed particularly strong performance, with a discovery AUC of 0.95 and a validation AUC of 0.85, again outperforming all published signatures with a similar number of transcripts (**Table S6**).

Finally, the performance of the new 10-transcript multinomial signature was evaluated, which enables simultaneous discrimination between VIR, BAC, and INF states. The multinomial signature performed similarly to the new binary signatures in both the discovery and validation cohorts for the 3 categories (**Table S6; Figure 6**). In the discovery cohort, the multinomial signature achieved strong class-wise performance, with AUCs of 0.94 for VIR, 0.89 for BAC, and 0.95 for INF classification. These values exceeded AUCs achieved by most of the previously published binary signatures comprising a larger number of transcripts and all AUCs obtained from signatures with a fewer number of transcripts, while simultaneously addressing a multi-class classification task (**Table S6**). In the validation cohort, the multinomial model retained robust performance across all classes (viral AUC=0.89, bacterial AUC=0.79, inflammatory AUC=0.85), demonstrating consistent generalizability (**Table S6; Figure 6**). Remarkably, in the validation cohort, the multinomial signature outperformed previously published binary signatures of comparable sizes in distinguishing bacterial and inflammatory samples.

### Multi-class signature in the confirmation cohort

To further assess the clinical applicability and cross-platform robustness of the multiclass signature, its performance was evaluated in an independent confirmation cohort using digital PCR (dPCR), a clinically deployable technology suitable for routine molecular diagnostics. The confirmation cohort was demographically comparable to the other cohorts and differed from the discovery cohort only in donor age within the viral category (**Table S1**). Gene expression levels of the ten transcripts were quantified across the three diagnostic categories, with *IFI27*, *IFIT3*, *CLC*, and *LY6E* showing the most pronounced disease-specific expression patterns (**Figure 7A**). Classification based on the predicted probabilities generated by the re-trained multinomial model demonstrated excellent diagnostic performance, achieving AUC values of 0.98 (95%CI: 0.95–1.00) for VIR infections, 0.91 (95%CI: 0.80–1.00) for BAC infections, and 0.85 (95%CI: 0.71–1.00) for INF diseases (**Figure 7B, 7C**). Notably, these results closely mirrored those obtained in the independent RNA-seq validation cohort, confirming the robustness, reproducibility, and transferability of the transcriptomic signature across patient cohorts and analytical platforms. The clearest separation of predicted class probabilities was observed for the VIR *vs*. REST comparison (*P*=1.4×10^−6^), further supporting the excellent performance of the multiclass classifier for viral infections.

**Figure 7.**
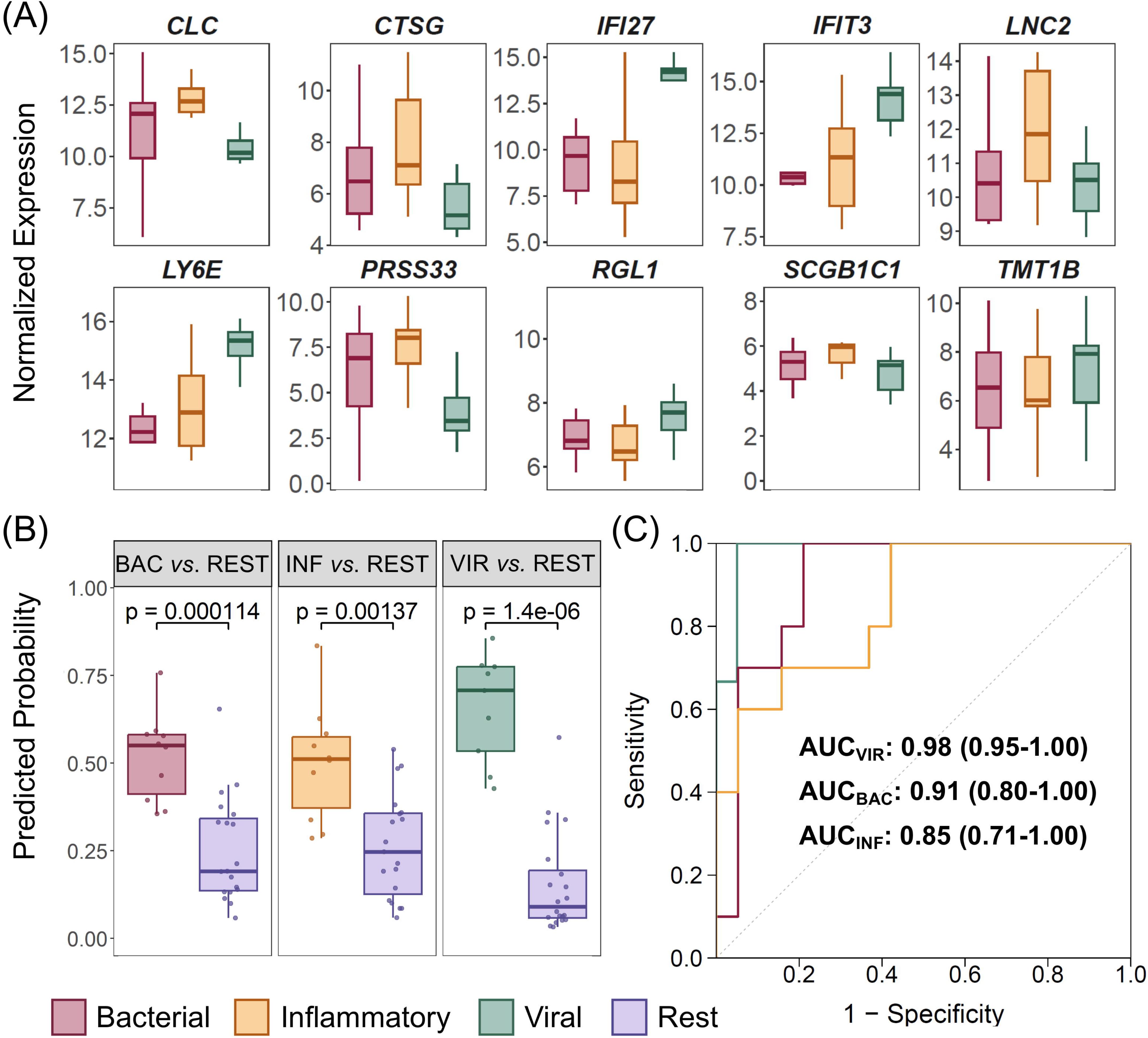
Independent validation of the multiclass transcriptomic signature using digital PCR (dPCR). (A) Distribution of normalized dPCR expression levels for the 10 transcripts included in the multiclass signature across the three diagnostic groups. (B) Predicted class probabilities for each one-versus-rest comparison. Statistical significance was assessed using two-sided Wilcoxon rank-sum tests. (C) Receiver operating characteristic (ROC) curves and corresponding area under the curve (AUC) values for the three one-versus-rest classification models.

To determine whether the signature could be reduced without compromising diagnostic performance, we subsequently evaluated a seven-gene model by excluding *SCGB1C1*, *RGL1*, and *TMT1B*, which showed the smallest differences in expression across diagnostic phenotypes in the dPCR confirmation cohort. The reduced model retained diagnostic performance comparable to that of the original ten-gene signature, with AUC values of 0.98 (95% CI: 0.95–1.00) for VIR infections, 0.90 (95% CI: 0.78–1.00) for BAC infections, and 0.85 (95% CI: 0.70–1.00) for INF diseases (**Figure S5**).

## Discussion

Accurate and timely discrimination between viral, bacterial, and non-infectious inflammatory diseases remains a major challenge in pediatric medicine ^2^, with important implications for patient management, antimicrobial stewardship, and healthcare resource utilization. The substantial clinical overlap among these conditions frequently hinders etiological diagnosis, leading to inappropriate treatment decisions. Unnecessary antibiotic exposure has been associated with long-term adverse health outcomes, including microbiota disruption, while accelerating antimicrobial resistance ^51–53^. Current diagnostic methods, including bacterial culture and molecular assays, are often too slow or insufficiently sensitive for early clinical decision-making ^54, 55^. Consequently, host-response biomarkers, particularly blood transcriptomic profiling, have emerged as promising tools for improving the etiological diagnosis of febrile illness by capturing disease-specific immune responses ^7, 10, 27, 31, 32, 56^.

Compared with conventional biomarkers (e.g., CRP and PCT), multi-gene transcriptomic signatures more comprehensively capture the host immune response and consistently achieve superior diagnostic performance ^57–61^. CRP lacks specificity because it is elevated in infectious and non-infectious inflammatory conditions, whereas the diagnostic utility of PCT is limited by suboptimal specificity, particularly in neonates, and by physiological increases during early life ^57, 59^.

Despite the growing number of published host transcriptomic signatures, few have progressed to clinically applicable diagnostic tests ^62^. This translational gap largely reflects limited reproducibility across independent cohorts and technological platforms, particularly for complex signatures that may be affected by overfitting and poor generalizability.

The present study constitutes the largest pediatric multi-cohort transcriptomic meta-analysis performed to date, integrating data from 2,450 children to derive and validate host-response signatures capable of discriminating viral, bacterial, and inflammatory diseases across independent populations and transcriptomic platforms. Signatures derived from a harmonized microarray discovery cohort were successfully validated in an independent RNA-seq meta-cohort and confirmed by dPCR, demonstrating reproducible performance across three independent transcriptomic technologies. Importantly, the inclusion of inflammatory diseases as a separate diagnostic category addresses an important limitation of many previous host-response classifiers, which focus exclusively on infectious diseases and therefore risk misclassifying inflammatory disorders ^63^.

A comprehensive comparative evaluation was performed to benchmark the newly derived transcriptomic signatures against a large collection of previously published models. The binary signatures consistently achieved high diagnostic accuracy while maintaining compact gene sets, with AUCs >0.88 in the training and test sets and >0.80 in the independent RNA-seq validation cohort. Within the discovery cohort, the inflammatory signature showed the highest accuracy (AUC=0.95 and 0.94 in the TA and TE sets, respectively), followed by the viral (AUC=0.93 in both sets) and bacterial (AUC=0.88 and 0.91, respectively) signatures. During external validation, the viral signature achieved the strongest performance (AUC=0.89). Compared with previously published models, the signatures ranked among the best-performing approaches while requiring substantially fewer transcripts. The bacterial and inflammatory signatures outperformed all published models with comparable or smaller gene sets, whereas the 8-transcript viral signature remained competitive with the best compact classifiers. Importantly, all binary signatures achieved AUCs above 0.75 across all age groups and above 0.81 in children younger than 24 months, the population at greatest risk of severe infection.

A compact 10-gene multinomial signature was also developed to simultaneously classify viral, bacterial, and inflammatory diseases. Despite addressing a more complex diagnostic task, it achieved performance comparable to the binary models, with AUCs >0.89 in the discovery cohort and >0.79 in the validation cohort. These results support its potential to guide clinical decision-making by simultaneously identifying patients requiring antibiotics, antiviral investigations or therapy, or alternative management for inflammatory diseases ^34, 64^. Performance remained consistently high across all age groups (AUC>0.78), with the best results observed in infants. Moreover, the multiclass signature matched or exceeded the performance of many published binary signatures despite requiring substantially fewer genes.

The multinomial signature retained its diagnostic accuracy in the dPCR confirmation cohort, achieving AUCs ranging from 0.85 to 0.98, comparable to those obtained in the discovery and validation cohorts. Its reproducible performance across independent cohorts and three transcriptomic technologies supports its robustness and clinical applicability. Together, these findings demonstrate that accurate multiclass diagnosis can be achieved using a single compact transcriptomic signature that outperforms previously reported multiclass classifiers in febrile children ^34^.

DCA provided a clinically relevant assessment of whether the improved discrimination achieved by the transcriptomic signatures translated into better clinical decision-making. Unlike conventional performance metrics, DCA quantifies clinical utility by incorporating decision thresholds and the consequences of diagnostic decisions ^65^. Although increasingly used in predictive modeling, DCA has rarely been applied to transcriptomic signatures for infectious diseases. Across both bacterial and viral infection scenarios, all transcriptomic models outperformed the default “treat-all” and “treat-none” strategies over a broad range of clinically relevant threshold probabilities, supporting their applicability across diverse clinical settings. Importantly, both the binary and multinomial signatures consistently demonstrated greater clinical utility than CRP, highlighting the limitations of single-biomarker approaches. For bacterial infections, the greatest net benefit was observed at higher decision thresholds, suggesting potential value for guiding antibiotic initiation. In contrast, viral classification showed greater benefit at lower thresholds, supporting early triage, additional diagnostic testing, or antiviral treatment while reducing unnecessary antibiotic use. The multinomial model maintained a consistently positive net benefit across the threshold range most relevant to clinical practice, outperforming CRP for both bacterial and viral classification. Whereas the clinical utility of CRP progressively declined and approached zero at higher thresholds, the transcriptomic signatures preserved a favorable balance between true-positive and false-positive decisions, demonstrating greater robustness for antibiotic-related decision-making. Overall, these findings highlight the added value of host transcriptomic profiling for guiding the differential diagnosis and management of infectious and inflammatory diseases beyond conventional biomarkers.

A major strength of this study is the derivation of diagnostic signatures from the largest harmonized pediatric multi-cohort transcriptomic dataset assembled to date, integrating multiple independent studies and transcriptomic technologies. This design minimizes study-specific biases, reduces overfitting, and improves generalizability across pathogens, clinical settings, and patient populations. Moreover, host-response signatures are inherently less affected than pathogen-centered diagnostic approaches by variables such as symptom duration, pathogen load, and prior antimicrobial exposure ^15^. The consistent performance observed across three independent transcriptomic platforms (microarray, RNA-seq, and dPCR) further demonstrates that the identified signatures reflect robust biological signals rather than platform-specific technical artifacts, supporting their translational potential. The reduction of the diagnostic signature from ten to seven genes should, however, be regarded as preliminary, given that it was evaluated in a relatively small dPCR confirmation cohort. Although the reduced model retained performance comparable to that of the original ten-gene signature, larger and prospectively designed validation cohorts will be necessary to establish whether this simplified signature provides consistent diagnostic performance or whether the original ten-gene model offers greater stability and accuracy when evaluated with greater statistical power. Moreover, additional prospective multicenter studies remain necessary to evaluate the clinical impact of the signatures, including turnaround time, cost-effectiveness, integration into existing diagnostic workflows, and performance in clinically challenging scenarios such as viral–bacterial coinfections. Further technological development will also be required to translate these signatures into rapid, robust, and scalable point-of-care assays suitable for routine clinical implementation.

Compact binary and multinomial blood transcriptomic signatures derived from harmonized multi-cohort datasets accurately distinguished viral, bacterial, and inflammatory diseases in children and consistently outperformed previously published host-response signatures. Their high diagnostic accuracy, reproducibility across independent cohorts and transcriptomic platforms, and demonstrated clinical utility support their potential as clinically deployable molecular biomarkers. Beyond improving etiological diagnosis, these findings reinforce host-response transcriptomics as a promising precision medicine strategy to optimize antimicrobial stewardship and clinical decision-making in pediatric infectious diseases.

## Data Availability

The gene expression data generated in this study and used in the confirmation cohort are publicly available in the Figshare repository (https://figshare.com/) under DOI: 10.6084/m9.figshare.33111542.

## Competing interests

The authors declare no competing interests.

## Consent for publication

All participants have given permission to the publication of the project’s findings.

## Ethics approval and consent to participate

Written informed consent was obtained from the legal guardians or responsible parties of all participating individuals in the confirmation cohort. The study was approved by the Ethics Committee of the Xunta de Galicia (GENDRES registration codes: 2010/015 2016/484; KawaTest registration code: 2022/320) and conducted in accordance with the principles outlined in the Declaration of Helsinki.

## Author’s contribution

ASE, AGC, and SVL conceived the project. AS and FMT provided funding. AGC, AS, and SVL carried out the analyses. AD, FMT, and IRC contributed to the clinical side of the project. AGC, AS, and SVL wrote the first draft of the manuscript. All authors critically reviewed the final version of the manuscript.

## Acknowledgements

This study received support by i) Instituto de Salud Carlos III (ISCIII): MANTRA-ID: PI25/00725; PNEUMO-ID360: DTS25/00083; TRINEO: PI22/00162; [to A.S.]; PNEU-MO-V-AI: PI25/01405; OMI-COVI-VAC: PI22/00406; [to F.M.-T.]; NIRSE-omic: PI24/00771 [to S.P.], and co-funded by the European Union; ii) GAIN: IN607B 2020/08 and IN607A 2023/02 [to A.S.]; GEN-COVID: IN845D 2020/23 and IIN607A2021/05 [to F.M.-T.] and IN677D 2024/06 [to A.G.-C.]; iii) ACIS: BI-BACVIR PRIS-3, CovidPhy SA304C, PneumoTrack PRIST-VAL [to A.S.] and Respisal PRIST-VAL [to F.M.-T.]; iv) Consorcio Centro de Investigación Biomédica en Red de Enfermedades Respiratorias (CB21/06/00103) [to F.M.-T. and A.S.]; and v) Spanish Ministry of Science and Innovation (MCIN)/Spanish Research Agency (AEI) and co-funded by the European Union: KAWA-TesT: PID2022-142156OB-I00 [to A.G.-C.]. AGC is supported by the Miguel Servet from ISCIII (CP23/00080) contract, funded by the Instituto de Salud Carlos III (ISCIII) and co-funded by the European Union. The funders were not involved in the study design, collection, analysis, interpretation of data, the writing of this article, or the decision to submit it for publication.

## Conflict of interests

Authors declare no conflicts of interests.

## Supplementary material

**Figure S1. PRISMA flow diagram describing the systematic search conducted to select (A) GEO (Gene Expression Omnibus) datasets and (B) PubMed publications related to transcriptomic signatures.**

**Figure S2. Individual normalization and batch correction of the selected publicly available microarray dataset.**

**Figure S3. PCA representation of the merging and batch-correction process applied to the different gene expression microarray datasets used in the meta-analysis (discovery cohort).**

**Figure S4. PCA representation of the merging and batch-correction process applied to the different RNA-seq datasets used in the meta-analysis (validation cohort).**

**Figure S5. Evaluation of the refined seven-gene multiclass transcriptomic signature derived from the original ten-gene panel using digital PCR (dPCR).** (A) Predicted class probabilities for each one-*versus*-rest comparison. Statistical significance was assessed using two-sided Wilcoxon rank-sum tests. (B) Receiver operating characteristic (ROC) curves and corresponding area under the curve (AUC) values for the three one-*versus*-rest classification models.

**Table S1. Description of the studies included in the discovery (microarray), validation (RNA-seq), and confirmation (digital PCR) cohorts. Demographic characteristics of each cohort are provided, together with intra-and inter-cohort statistical comparisons for age and sex.** HC, healthy controls; KD, Kawasaki disease; HSP, Henoch–Schönlein purpura; JIA, juvenile idiopathic arthritis. Wilcoxon rank-sum and Fisher’s exact test were used to assess statistical significance between patient groups.

**Table S2. TaqMan expression assay IDs and reaction combinations used for the dPCR 10-transcript signature validation in the confirmation cohort. Table S3. Gene list of previously published transcriptomic signatures considered in the present study. HGNC gene symbol list used for the comparison analysis.**

**Table S4. Differentially expressed genes (DEGs) candidates derived from the comparisons VIR *vs*. REST (BAC+INF), BAC *vs*. REST (VIR+INF), and INF *vs*. RES (VIR+BAC), meeting the established criteria (*P*_adj_<0.05, |log_2_FC)|>1, average log_2_ expression >4).** VIR: Viral; BAC: Bacterial; INF: Inflammatory.

**Table S5. Model coefficients for the genes included in the binary and multi-class signatures.**

**Table S6. Performance of previously published transcriptomic signatures in infectious diseases compared to the signatures derived from the present study in the discovery (in the complete discovery cohort and in the training [TA] and test [TE] sets separately) and validation cohorts.** BAC: Bacterial infection; VIR: Viral infection; HC: Healthy control; INF: Inflammatory diseases.

**Supplementary Text 1**. Consortia membership.

## Notes

### Competing Interest Statement

The authors have declared no competing interest.

