## Supplementary figures and images for "Host gene-expression signatures accurately distinguish bacterial, viral, and inflammatory diseases in febrile children across multiple cohorts"

### Figure S1

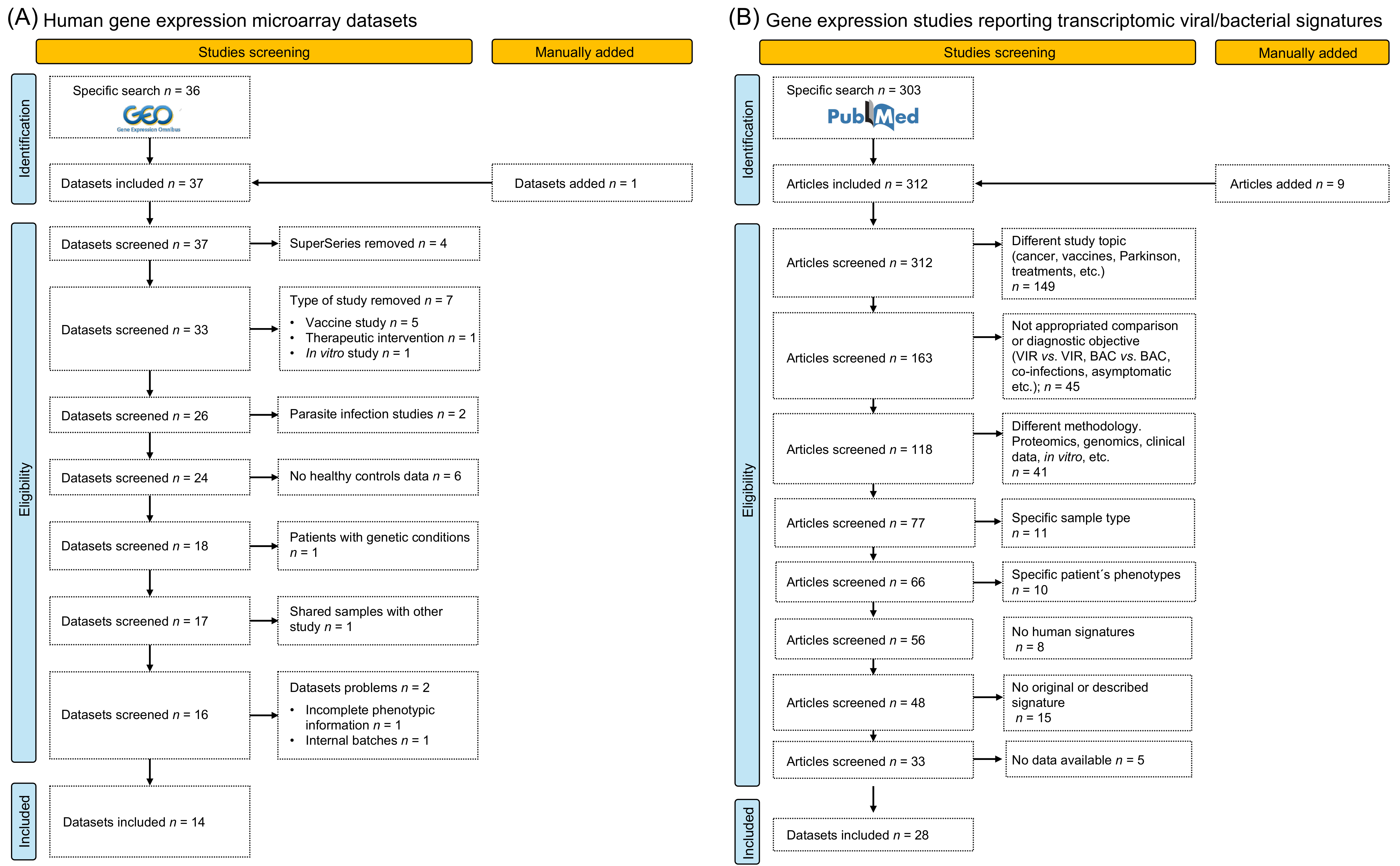

### Figure S2

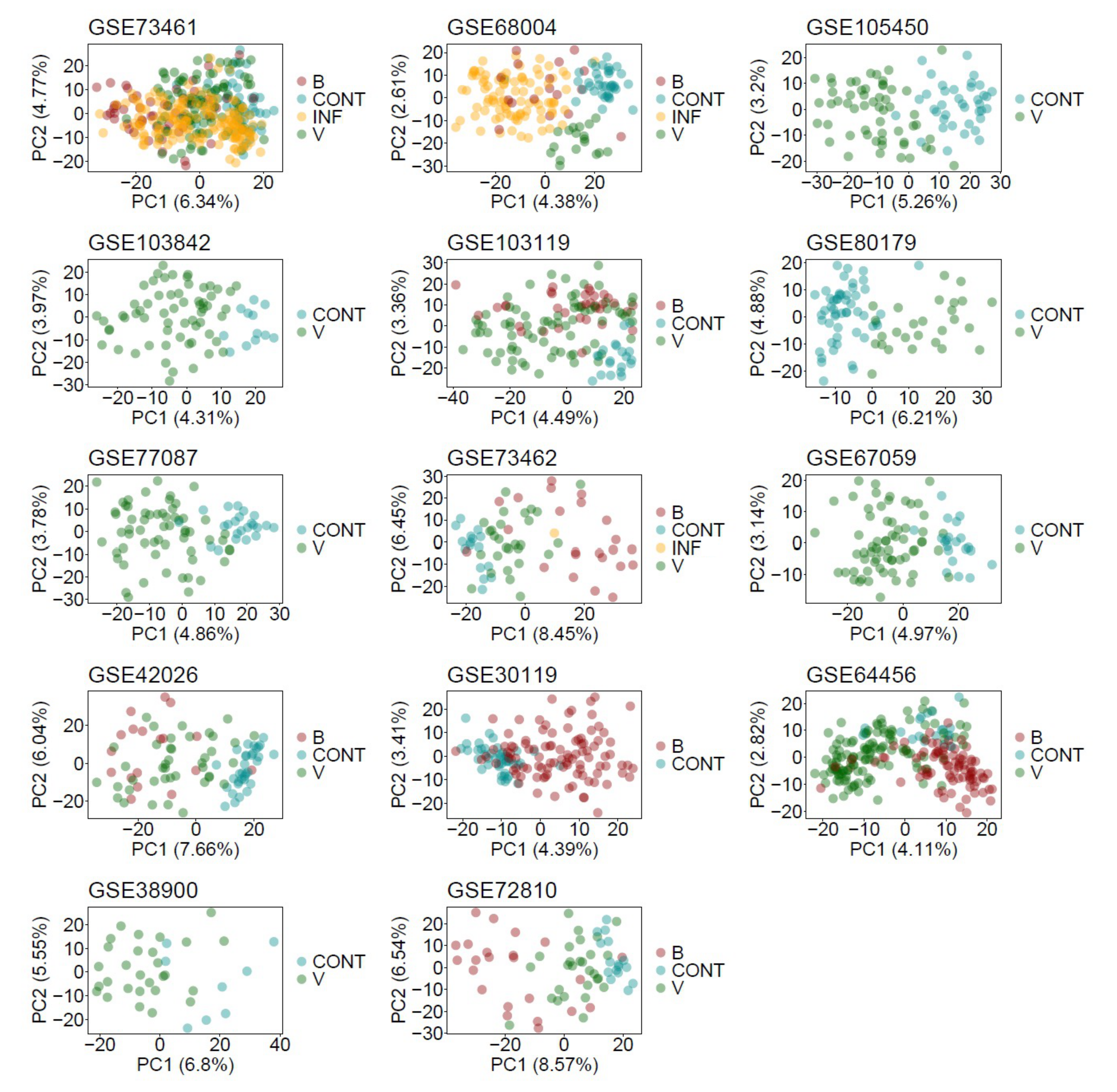

### Figure S3

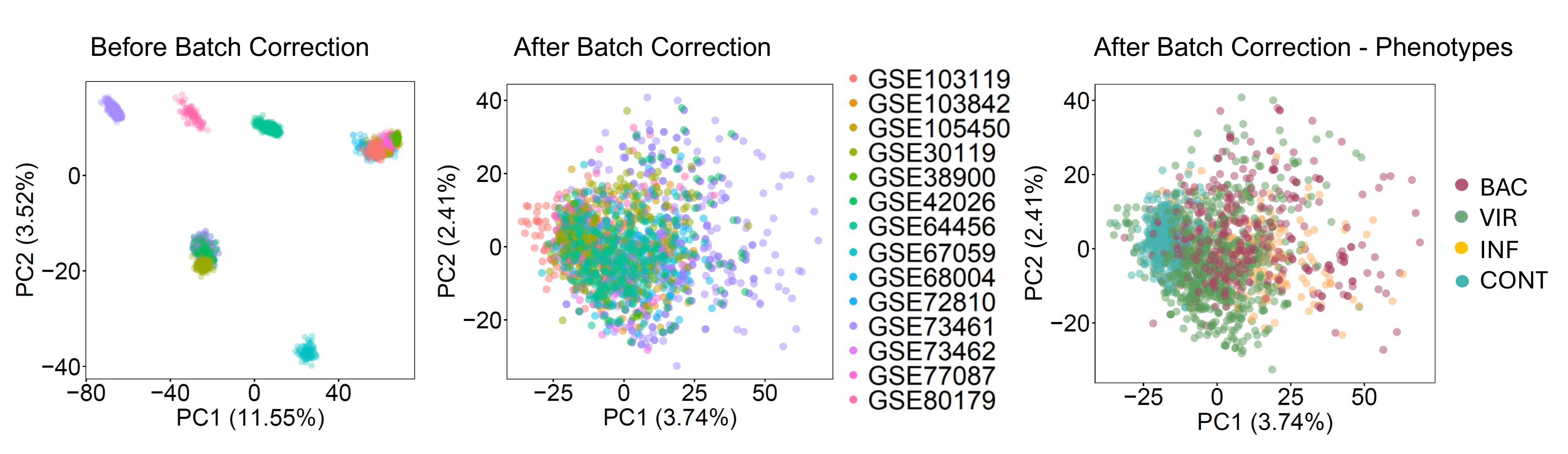

### Figure S4

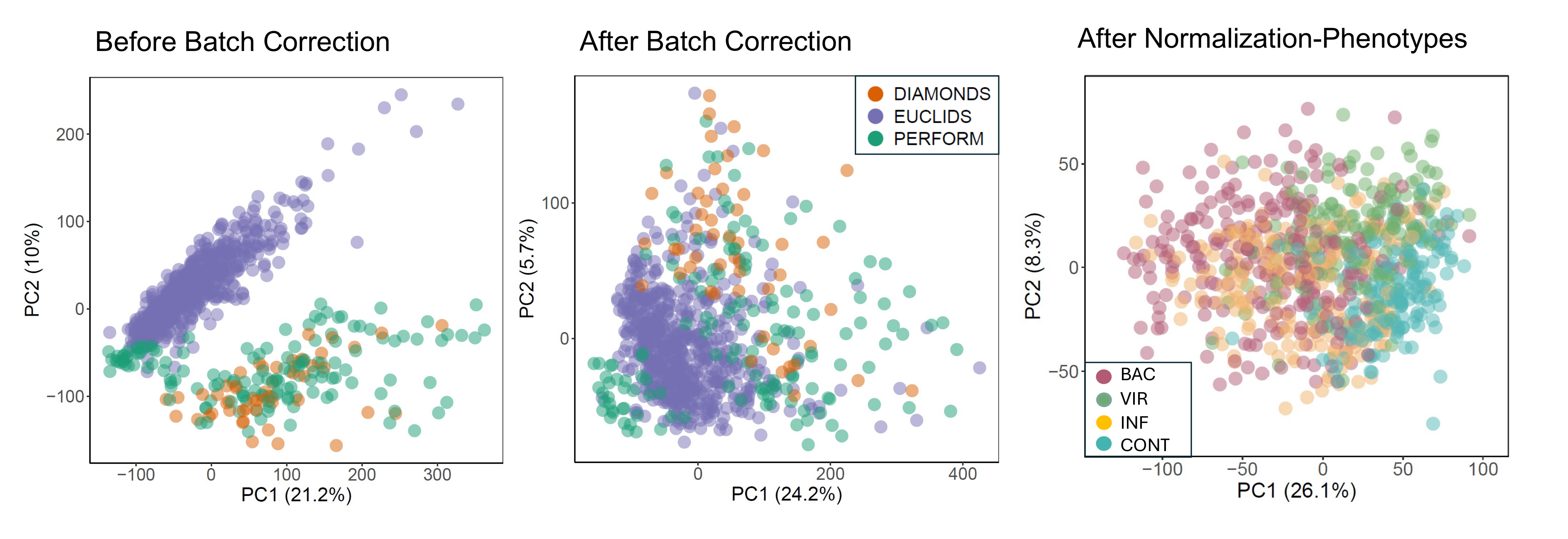

### Figure S5

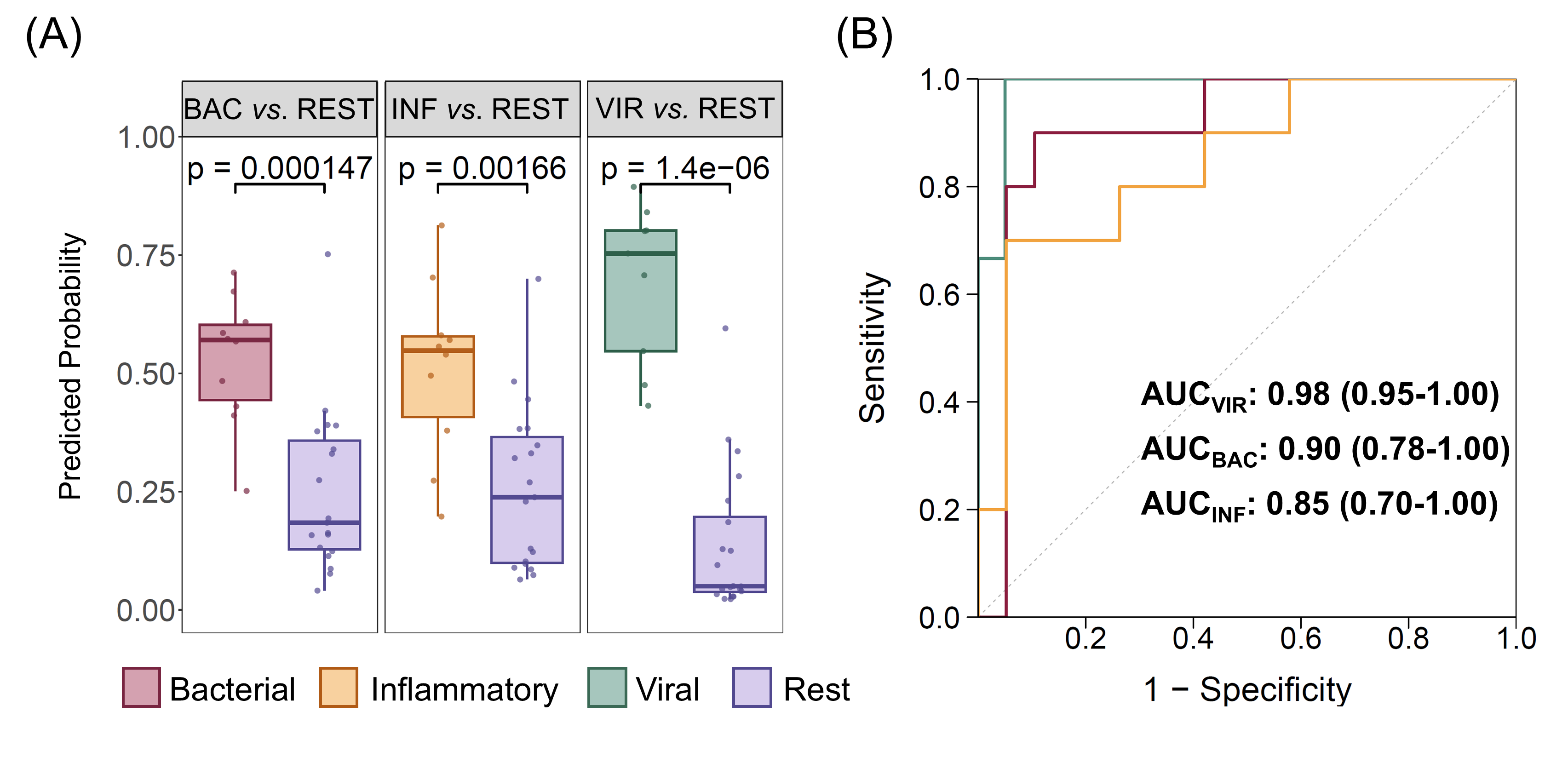
