## Supplementary Text 1 for "Host gene-expression signatures accurately distinguish bacterial, viral, and inflammatory diseases in febrile children across multiple cohorts"

**Consortia**

### **EUCLIDS consortium**

Michael Levin, Lachlan Coin, Stuart Gormley, Shea Hamilton, Jethro Herberg, Bernardo Hourmat, Clive Hoggart, Myrsini Kaforou, Vanessa Sancho-Shimizu, Victoria Wright, Amina Abdulla, Paul Agapow, Maeve Bartlett, Evangelos Bellos, Hariklia Eleftherohorinou, Rachel Galassini, David Inwald, Meg Mashbat, Stefanie Menikou, Sobia Mustafa, Simon Nadel, Rahmeen Rahman, Clare Thakker, Sumit Bokhandi, Sue Power, Heather Barham, Nazima Pathan, Jenna Ridout, Deborah White, Sarah Thurston, Saul Faust, Sanjay Patel, Jenni McCorkell, Patrick Davies, Lindsey Crate, Helen Navarra, Stephanie Carter, Ramesh Ramaiah, Rekha Patel, Catherine Tuffrey, Andrew Gribbin, Sharon McCready, Mark Peters, Katie Hardy, Fran Standing, Lauran O’Neill, Eugenia Abelake, Akash Deep, Eniola Nsirim, Andrew Pollard, Louise Willis, Zoe Young, C. Royad, Sonia White, PM. Fortune, Phil Hudnott, Federico Martinón Torres, Antonio Salas Ellacuriaga, Fernando Álvez González, Ruth Barral-Arca, Miriam Cebey-López, María José Curras-Tuala, Natalia García, Luisa García Vicente, Alberto Gómez-Carballa, Jose Gómez Rial, Andrea Grela Beiroa, Antonio Justicia Grande, Pilar Leboráns Iglesias, Alba Elena Martínez Santos, Nazareth Martinón-Torres, José María Martinón Sánchez, Beatriz Morillo Gutiérrez, Belén Mosquera Pérez, Pablo Obando Pacheco, Jacobo Pardo-Seco, Sara Pischedda, Sandra Viz Lasheras, Irene Rivero Calle, Carmen Rodríguez-Tenreiro, Lorenzo Redondo-Collazo, Sonia Serén Fernández, María del Sol Porto Silva, Ana Vega, Jose Manuel Fernández García, María Elena Gamborino Caramés, María Sol Rodríguez Calvo, Marta Aldonza Torres, Vanesa Álvarez Iglesias, Carmen Curros Novo, Ramón, Isabel Rego Lijo, Ana Isabel Dacosta Urbieta, Wiktor Dominik Nowak, Miriam Taboada Puga, Lucía Vilanova Trillo, Susana Beatriz Reyes, María Cruz León León, Álvaro Navarro Mingorance, Xavier Gabaldó Barrios, Eider Oñate Vergara, Andrés Concha Torre, Ana Vivanco, Reyes Fernández, Francisco Giménez Sánchez, Miguel Sánchez Forte, Pablo Rojo, J. Ruiz Contreras, Alba Palacios, Cristina Epalza Ibarrondo, Elizabeth Fernandez Cooke, Marisa Navarro, Cristina Álvarez Álvarez, María José Lozano, Eduardo Carreras, Sonia Brió Sanagustín, Olaf Neth, Mª del Carmen Martínez Padilla, Luis Manuel Prieto Tato, Sara Guillén, Laura Fernández Silveira, David Moreno, Ronald de Groot, A. Marceline Tutu van Furth, Michiel van der Flier, Navin P. Boeddha, Gertjan J. A. Driessen, Jan A. Hazelzet, Taco W. Kuijpers, Dasja Pajkrt, Elisabeth A. M. Sanders, Diederik van de Beek, A. van der Ende, H.L.A. Philipsen, A.O.A. Adeel, M.A. Breukels, D.M.C. Brinkman, C.C.M.M. de Korte, E. de Vries, W.J. de Waal, R. Dekkers, A. Dings-Lammertink, R.A. Doedens, A.E. Donker, M. Dousma, T.E. Faber, G.P.J.M. Gerrits, J.A.M. Gerver, J. Heidema, J. Homan-van der Veen, M.A.M. Jacobs, N.J.G. Jansen, P. Kawczynski, K. Klucovska, M.C.J. Kneyber, Y. Koopman-Keemink, V.J. Langenhorst, J. Leusink, B.F. Loza, I.T. Merth, C.J. Miedema, C. Neeleman, J.G. Noordzij, C.C. Obihara, A.L.T. van Overbeek – van Gils, G.H. Poortman, S.T. Potgieter, J. Potjewijd, P.P.R. Rosias, T. Sprong, G.W. ten Tussher, B.J. Thio, G.A. Tramper-Stranders, M. van Deuren, H. van der Meer, A.J.M. van Kuppevelt, A.M. van Wermeskerken, W.A. Verwijs, T.F.W. Wolfs, Luregn J Schlapbach, Philipp Agyeman, Christoph Aebi, Christoph Berger, Eric Giannoni, Martin Stocker, Klara M Posfay-Barbe, Ulrich Heininger, Sara Bernhard-Stirnemann, Anita Niederer-Loher, Christian Kahlert, Paul Hasters, Christa Relly, Walter Baer, Enitan Carrol, Stéphane Paulus, Hannah Frederick, Rebecca Jennings, Joanne Johnston, Rhian Kenwright, Colin G Fink, Elli Pinnock, Marieke Emonts, Emma Lim, Lucille Valentine, Karen Allen, Kathryn Bell, Adora Chan, Stephen Crulley, Kirsty Devine, Daniel Fabian, Sharon King, Paul McAlinden, Sam McDonald, Anne McDonnell, Ailsa Pickering, Evelyn Thomson, Amanda Wood, Diane Wallia. Rachel Agbeko, Suzanne Anderson, Fatou Secka, Kalifa Bojang, Isatou Sarr, Ngane Kebbeh, Gibbi Sey, Momodou Saidykhan, Fatoumatta Cole, Gilleh Thomas, Martin Antonio, Werner Zenz, Daniela S. Klobassa, Alexander Binder, Nina A. Schweintzger, Manfred Sagmeister, Hinrich Baumgart, Markus Baumgartner, Uta Behrends, Ariane Biebl, Robert Birnbacher, Jan-Gerd Blanke, Carsten Boelke, Kai Breuling, Jürgen Brunner, Maria Buller, Peter Dahlem, Beate Dietrich, Ernst Eber, Johannes Elias, Josef Emhofer, Rosa Etschmaier, Sebastian Farr, Ylenia Girtler, Irina Grigorow, Konrad Heimann, Ulrike Ihm, Zdenek Jaros, Hermann Kalhoff, Wilhelm Kaulfersch, Christoph Kemen, Nina Klocker, Bernhard Köster, Benno Kohlmaier, Eleni Komini, Lydia Kramer, Antje Neubert, Daniel Ortner, Lydia Pescollderungg, Klaus Pfurtscheller, Karl Reiter, Goran Ristic, Siegfried Rödl, Andrea Sellner, Astrid Sonnleitner, Matthias Sperl, Wolfgang Stelzl, Holger Till, Andreas Trobisch, Anne Vierzig, Ulrich Vogel, Christina Weingarten, Stefanie Welke, Andreas Wimmer, Uwe Wintergerst, Daniel Wüller, Andrew Zaunschirm, Ieva Ziuraite, Veslava Žukovskaja.

**PERFORM and DIAMONDS consortiums**

Michael Levin, Aubrey Cunnington, Jethro Herberg, Myrsini Kaforou, Victoria J Wright, Evangelos Bellos, Claire Broderick, Samuel Channon-Wells, Samantha Cooray, Tisham De, Giselle D'Souza, Amedine Duret, Anikta Duseja, Leire Estamiana Ellorrieta, Diego Estrada-Rivadeneyra, Rachel Gallassini, Dominic Habgood-Coote, Shea Hamilton, Heather Jackson, James Kavanagh, Ilana Keren, Mahdi Moradi Marjeneh, Stephanie Menikou, Samuel Nichols, Ruud Nijman, Harsita Patel, Ivana Pennisi, Oliver Powell, Ruth Reid, Priyen Shah, Ortensia Vito, Elizabeth Whittaker, Clare Wilson, Rebecca Womersley, Amina Abdulla, Sarah Darnell, Sobia Mustafa, Pantelis Georgiou, Jesus Rodriguez-Manzano, Nicolas Moser, Michael Carter, Paul Wellman, Shane Tibby, Jonathan Cohen, Francesca Davis, Julia Kenny, Marie White, Matthew Fish, Aislinn Jennings, Manu Shankar-Hari, Katy Fidler, Dan Agranoff, Viven Richmond, Matthew Seal, Saul Faust, Dan Owen, Ruth Ensom, Sarah McKay, Diana Mondo, Mariya Shaji, Rachel Schranz, Prita Rughani, Amutha Anpananthar, Susan Liebeschuetz, Anna Riddell, Divya Divakaran, Louise Han, Nosheen Khalid, Ivone Lancoma-Malcolm, Jessica Schofield, Teresa Simagan, Mark Peters, Alasdair Bamford, Lauran O'Neill, Nazima Pathan, Esther Daubney, Deborah White, Melissa Heightman, Sarah Eisen, Terry Segal, Lucy Wellings, Simon B Drysdale, Nicole Branch, Lisa Hamzah, Heather Jarman, Maggie Nyirenda, Lisa Capozzi, Emma Gardiner, Robert Moots, Madga Nasher, Anita Hanson, Michelle Linforth, Sean O'Riordan, Donna Ellis, Akash Deep, Ivan Caro, Fiona Shackley, Arianna Bellini, Stuart Gormley, Samira Neshat, Barnaby J Scholefield, Ceri Robbins, Helen Winmill, Stéphane C Paulus, Andrew J Pollard, Mark Anthony, Sarah Hopton, Danielle Miller, Zoe Oliver, Sally Beer, Bryony Ward, Shrijana Shrestha, Meeru Gurung, Puja Amatya, Bhishma Pokhrel, Sanjeev Man Bijukchhe, Madhav Chandra Gautam, Sarah Kelly, Peter O'Reilly, Sonu Shrestha, Federico Martinón-Torres, Antonio Salas, Fernando Álvez González, Sonia Ares Gómez, Xabier Bellos, Mirian Ben García, Fernando Caamaño Viña, Sandra Carnota, María José Curras-Tuala, Ana Dacosta Urbieta, Carlos Durán Suárez, Isabel Ferreiros Vidal, Luisa García Vicente, Alberto Gómez-Carballa, Jose Gómez Rial, Pilar Leboráns Iglesias, Narmeen Mallah, Nazareth Martinón-Torres, José María Martinón Sánchez, Belén Mosquera Pérez, Jacobo Pardo-Seco, Sara Pischedda, Sara Rey Vázquez, Sandra Viz Lasheras, Alba Camino Mera, Lúa Castelo Martínez, Nour El Zaharaa Mallah, Patricia Regueiro Casuso, Irene Rivero Calle, Carmen Rodríguez-Tenreiro, Lorenzo Redondo-Collazo, Sonia Serén Fernández, Marisol Vilas Iglesias, Enitan D Carrol, Elizabeth Cocklin, Rebecca Beckley, Abbey Bracken, Ceri Evans, Aakash Khanijau, Rebecca Lenihan, Nadia Lewis-Burke, Karen Newall, Sam Romaine, Jennifer Whitbread, Maria Tsolia, Irini Eleftheriou, Nikos Spyridis, Maria Tambouratzi, Despoina Maritsi, Antonios Marmarinos, Marietta Xagorari, Lourida Panagiota, Pefanis Aggelos, Akinosoglou Karolina, Gogos Charalambos, Maragos Markos, Voulgarelis Michalis, Stergiou Ioanna, Marieke Emonts, Emma Lim, John Isaacs, Kathryn Bell, Stephen Crulley, Daniel Fabian, Evelyn Thomson, Diane Wallia, Caroline Miller, Ashley Bell, Mathew Rhodes, Fabian J S Van der Velden, Geoff Shenton, Ashley Price, Owen Treloar, Daisy Thomas, Pablo Rojo, Cristina Epalza, Serena Villaverde, Sonia Márquez, Manuel Gijón, Romina Varchetta, Fátima Machín, Laura Cabello, Irene Hernández, Lourdes Gutiérrez, Ángela Manzanares, T W Taco Kuijpers, M Martijn Van de Kuip, A M Marceline Van Furth, J M Merlijn Van den Berg, Giske Biesbroek, Floris Verkuil, Carlijn C W Van der Zee, Dasja Pajkrt, Michael Boele van Hensbroek, Dieneke Schonenberg, Mariken Gruppen, Sietse Nagelkerke, Machiel H Jansen, Ines Goetschalckx, Lorenza Romani, Maia De Luca, Sara Chiurchiù, Costanza Tripiciano, Stefania Mercadante, Clementien L Vermont, Henriëtte A Moll, Dorine M Borensztajn, Nienke N Hagedoorn, Chantal Tan, Joany Zachariasse, W Dik, Ching-Fen Shen, Dace Zavadska, Sniedze Laivacuma, Aleksandra Rudzate, Diana Stoldere, Arta Barzdina, Elza Barzdina, Monta Madelane, Dagne Gravele, Dace Svile, Romain Basmaci, Noémie Lachaume, Pauline Bories, Raja Ben Tkhayat, Laura Chériaux, Juraté Davoust, Kim-Thanh Ong, Marie Cotillon, Thibault de Groc, Sébastien Le, Nathalie Vergnault, Hélène Sée, Laure Cohen, Alice de Tugny, Nevena Danekova, Marine Mommert-Tripon, Karen Brengel-Pesce, Marko Pokorn, Mojca Kolnik, Tadej Avčin, Tanja Avramoska, Natalija Bahovec, Petra Bogovič, Lidija Kitanovski, Mirijam Nahtigal, Lea Papst, Tina Plankar Srovin, Frac Strle, Katarina Vincek, Michiel van der Flier, Wim J E Tissing, Rosalie M Wösten-van Asperen, Sebastiaan J Vastert, Daniel C Vijlbrief, Louis J Bont, Coco R Beudeker, Philipp Agyeman, Christoph Aebi, Nina Schöbi, Mariama Usman, Stefanie Schlüchter, Luregn Schlapbach, Cornelia Hagmann, Florian Zapf, Philipp Baumann, Barbara Brotschi, Elisa Zimmermann, Marion Meier, Kathrin Weber, Colin Frink, Marie Voice, Leo Calvo-Bado, Michael Steele, Jennifer Holden, Andrew Taylor, Ronan Calvez, Catherine Davies, Benjamin Evans, Jake Stevens, Peter Matthews, Kyle Billing, Werner Zenz, Alexander Binder, Benno Kohlmaier, Daniela S Kohlfürst, Nina A Schweintzge, Christoph Zurl, Susanne Hösele, Piyush G Gampawar, Barbara Kapo, Manuel Leitner, Lena Pölz, Alexandra Rusu, Glorija Rajic, Bianca Stoiser, Martina Strempfl, Manfred G Sagmeister, Sebastian Bauchinger, Martin Benesch, Astrid Ceolotto, Ernst Eber, Siegfried Gallistl, Harald Haidi, Almuthe Hauer, Christa Hude, Andrea Kapper, Markus Keldorfer, Sabine Löffler, Tobias Niedrist, Heidemarie Pilch, Andreas Pfleger, Klaus Pfurtscheller, Siegfried Rödl, Andrea Skrabi-Baumgartner, Volker Strenger, Elmar Wallner, Maike K Tauchert, Ulrich von Both, Laura Kolberg, Patricia Schmied, Ioanna Mavridi, Irene Alba-Alejandre, Katharina Danhauser, Nikolaus Haas, Matthias Griese, Tobias Feuchtinger, Sabrina Juranek, Matthias Kappler, Eberhard Lurz, Esther Maier, Karl Reiter, Carola Schoen, Sebastian Schroepf, Shunmay Yeung, Manuel Dewez, David Bath, Elizabeth Fitchett, Fiona Cresswell, Effua Usuf, Kalifa Bojang, Anna Roca, Isatou Sarr, Momodou Saidykhan, Ebrahim Ndure, Pedro Madrigal, Silvie Fexova, Artur Sulik, Kacper Toczylowski, Dawid Lewandowski, Victoria Wright, Lucas Baumard, Rachel Galassini, Clive Hoggart, Sara Hourmat, Ian Maconochie, Naomi Lin, Ivonne Pena Pas, Hannah Shailes, Ladan Ali, Rikke Jorgensen, Salina Persand, Molly Stevens, Eunjung Kim, Benjamin Pierce, Julia Dudley, Vivien Richmond, Emma Tavliavini, Ching-Chuang Liu, Shih-Min Wang, Cristina Balo Farto, Ruth Barral-Arca, Maria Barreiro Castro, Xabier Bello, Miriam Cebey-López, Lidia Piñeiro Rodríguez, Miguel Sadiki Ora, Cristina Serén Trasorras, Dace Zavadaska, Anda Balode, Dārta Deksne, Dace Gardovska, Ilze Grope, Anija Meiere, Ieva Nokalna, Jana Pavāre, Zanda Pučuka, Katrīna Selecka, Aleksandra Sidorova, Urzula Nora Urbāne, Syed M A Zaman, Fatou Secka, Suzanne Anderson, Saffiatou Darboe, Samba Ceesay, Umberto D'alessandro, Luregn J Schlapbach, Verena Wyss, Eric Giannoni, Martin Stocker, Klara M Posfay-Barbe, Ulrich Heininger, Sara Rey Bernhard-Stirnemann, Anita Niederer-Loher, Christian Kahlert, Giancarlo Natalucci, Christa Relly, Thomas Riedel, Christoph Berger, Stéphane Paulus, Rebecca Jennings, Joanne Johnston, Simon Leigh, Antonis Marmarinos, Kelly Syggelou, Colin Fink, Nina A Schweintzger, Hinrich Baumgart, Gunther Gores, Harald Haidl, Larissa Krenn, Gudrun Nordberg, Andrea Skrabl-Baumgartner, Matthias Sperl, Laura Stampfer, Holger Till, Andreas Trobisch, Juan Emmanuel Dewez, Martin Hibbered, Alec Miners, Catherine Wedderburn, Anne Meierford, Baptiste Leurent, Ronald de Groot, Marien I de Jonge, Koen van Aerde, Wynard Alkema, Bryan van den Broek, Jolein Gloerich, Alain J van Gool, Stefanie Henriet, Martijn Huijnen, Ria Philipsen, Esther Willems, G P J M Gerrits, M van Leur, J Heidema, L de Haan, C J Miedema, C Neeleman, C C Obihara, G A Tramper-Stranders, Rama Kandasamy, Michael J Carter, Daniel O'Connor, Sagida Bibi, Dominic F Kelly, Stephen Thorson, Imran Ansari, David R Murdoch, Lucille Valentine, Karen Allen, Adora Chan, Kirsty Devine, Sharon King, Paul McAlinden, Sam McDonald, Anne McDonnell, Ailsa Pickering, Amanda Wood, Phil Woodsford, Frances Baxter, Matthew Rhodes, Rachel Agbeko, Christine Mackerness, Bryan Baas, Lieke Kloosterhuis, Wilma Oosthoek, Tasnim Arif, Joshua Bennet, Kalvin Collings, Ilona van der Giessen, Alex Martin, Aqeela Rashid, Emily Rowlands, Gabriella de Vries, Fabian van der Velden, Joshua Soon, Manuela Zwerenz, Judith Bruschbeck, Christoph Bidlingmaier, Vera Binder, Julia Keil, Georg Muench, François Mallet, Alexandre Pachot, Marine Mommert, Petra Prunk, Veronika Osterman, Taco Kuijpers, Ilse Jongerius, J M van den Berg, D Schonenberg, A M Barendregt, D Pajkrt, M van der Kuip, A M van Furth, Evelien Sprenkeler, Judith Zandstra, G van Mierlo, Judy Geissler.

**GENDRES network**

Miriam Cebey López, Antonio Salas Ellacuriaga, Ana Vega Gliemmo, José Peña Guitián, Alexa Regueiro, Antonio Justicia Grande, Leticia Pías Peleteiro, María López Sousa, María Jose de Castro, Carmen Curros Novo, Elena Rodrigo, Miriam Puente Puig, Rosaura Leis Trabazo, Nazareth Martinón Torres, Alberto Gómez Carballa, Jacobo Pardo Seco, Sara Pischedda, José María Martinón Sánchez, Belén Mosquera Pérez, Isabel Villanueva González, Lorenzo Redondo Collazo, Carmen Rodríguez-Tenreiro, María del Sol Porto Silva and Federico Martinón Torres (Área Asistencial Integrada de Pediatría and GENVIP, Hospital Clínico Universitario, Santiago de Compostela); Máximo Francisco Fraga Rodríguez, Orlando Fernández Lago, José Ramón Antúnez (Biobank, Servicio Anatomía Patológica, Hospital Clínico Universitario, Santiago de Compostela); Enrique Bernaola Iturbe, Laura Moreno Galarraga, Jorge Álvarez, Mercedes Herranz, Francisco Gil, Eva Gembero, Jorge Rodríguez (Hospital Materno Infantil Virgen del Camino, Pamplona); Teresa González López, Delfina Suarez Vázquez, Ángela Vázquez Vázquez, Susana Rey García, Nathalie Carreira Sande, Ana López Fernández, Nuria Romero Pérez (Complejo Hospitalario Universitario de Orense); José Antonio Couceiro Gianzo, Nazareth Fuentes Perez (Complejo Hospitalario Universitario de Pontevedra); Francisco Giménez Sánchez, Miguel Sánchez Forte (Hospital Torrecárdenas, Almería); Cristina Calvo Rey, María Luz García García, Iciar Olabarrieta Arnal, Adelaida Fernández Rincón (Hospital Severo Ochoa de Madrid); Ignacio Oulego Erroz, David Naranjo Vivas, Santiago Lapeña, Paula Alonso Quintela, Jorge Martínez Sáenz de Jubera, Estibaliz Garrido García (Hospital de León); Ana Grande Tejada (Hospital Materno Infantil de Badajoz); Cristina Calvo Monge, Eider Oñate Vergara (Hospital de Donostia, San Sebastián); Jesús de la Cruz Moreno, Mª Carmen Martínez Padilla, Beatriz Jiménez Jurado, Carmen Santiago Gutiérrez, María Esther Vidaurreta del Castillo (Complejo Hospitalario de Jaén); Manuel Baca Cots (Hospital Quirón, Málaga), David Moreno Pérez, Ana Cordón Martínez, Antonio Urda Cardona, José Miguel Ramos Fernández, Esmeralda Núñez Cuadros (Hospital Carlos Haya, Málaga); Susana Beatriz Reyes, María Cruz León León, Santiago Alfayate (Hospital Virgen de la Arrixaca, Murcia); Cristina Calvo, Carlos Grasa, Cristian Quintana Ortega, Leticia La Banda Montalvo, María Lopez Cerdán, Ana Dominguez Castells (Hospital Universitario La Paz, Madrid); Francisco Giménez Sánchez (Hospital Inmaculada); Andrés J. Alcaraz Romero, Diego Bautista Lozano, Sara Uillen Martin (Hospital Universitario de Getafe); Roi Piñeiro (Hospital General de Villalba); Juan Ignacio Sánchez Díaz, Alba Palacios Cuesta (Hospital 12 Octubre); Elvira González Salas, Sira Fernández De Miguel (Hospital Clínico de Salamanca); Belen Joyanes Abancens, Esther Aleo Lujan (Hospital clínico San Carlos); Alfredo Tagarro García, María Luisa Herreros, Rut del Valle, Libertad Latorre Navarro (Hospital Universitario Infanta Sofía); María Concepción Zazo Sanchidrián, Mariano Esteban, Marta González Lorenzo, Mª Carmen Vicent Castello (Hospital General Universitario de Alicante); Lorena Moreno Requena, Juan Luis Santos Pérez (Complejo Hospitalario Universitario de Granada); César Gavilán Martín, Lucía González-Moro Azorín (Hospital Universitario de San Juan de Alicante); Monterrat López Franco, Manuel Silveira Cancela (Hospital de Burela); María José Cilleruelo Ortega, Luz Golmayo, Francisca Portero Azorín (Hospital Universitario Puerta de Hierro-Majadahonda); Andrés Concha Torre, Lucía Rodríguez García (Hospital Universitario Central de Asturias); Carlos Rodrigo Gonzalo de Liria, Andrés Antón Pagarolas (Hospital Vall d’Hebron); Maria Méndez, Cristina Prat (Hospital Germans Trias i Pujol); Jesús López-Herce, Miriam García Samprudencio, Gema Manrique Martín, Paula García Casas, Débora Sanz Álvarez (Hospital General Universitario Gregorio Marañón); María Jesús Cabero (Hospital Valdecilla); Miguel Lillo Lillo, Marta Pareja (Hospital General de Albacete); Pablo Rojo, Cristina Epalza (Hospital 12 Octubre); Adriana Navas Carretero, Estefanía Barral, Miriam Herrera (Hospital Infanta Leonor); Elvira Cobo Vázquez (Hospital Universitario Fundación de Alcorcón); Elena del Castillo Navío (Hospital Materno Infantil de Badajoz); Patricia Flores Perez (Hospital del Niño Jesús); Paula García Casas (Hospital Ramón y Cajal); Esteban Gómez Sánchez, Juan Valencia Ramos (Hospital Universitario de Burgos); Francisco Javier Pilar Orive, Elva Rodríguez Merino (Hospital Universitario Cruces); Ana Pérez Aragón (Hospital Universitario Virgen de las Nieves de Granada); Mª Yolanda Ruiz del Prado (Hospital San Pedro); David Moreno, Beatriz Carazo (Hospital Carlos Haya); Jordi Antón (Hospital Sant Joan de Déu); María teresa Rives Ferreiro (Hospital Universitario de Navarra). Further details may be consulted at http://www.gendres.org, and www.genvip.eu.
